# Excess neuronal branching allows for innervation of specific dendritic compartments in cortex

**DOI:** 10.1101/529875

**Authors:** A D Bird, L H Deters, H Cuntz

**Affiliations:** Frankfurt Institute for Advanced Studies, Frankfurt-am-Main, 60438, Germany; Ernst Strüngmann Institute (ESI) for Neuroscience in cooperation with the Max Planck Society, Frankfurt-am-Main, 60528, Germany

**Keywords:** Synaptic contacts, Minimum spanning tree, Morphology, Connectome, Dendritic compartments

## Abstract

The connectivity of cortical microcircuits is a major determinant of brain function; defining how activity propagates between different cell types is key to scaling our understanding of individual neuronal behaviour to encompass functional networks. Furthermore, the integration of synaptic currents within a dendrite depends on the spatial organisation of inputs, both excitatory and inhibitory. We identify a simple equation to estimate the number of potential anatomical contacts between neurons; finding a linear increase in potential connectivity with cable length and maximum spine length, and a decrease with overlapping volume. This enables us to predict the mean number of candidate synapses for reconstructed cells, including those realistically arranged. We identify an excess of putative connections in cortical data, with densities of neurite higher than is necessary to reliably ensure the possible implementation of any given connection. We show that potential contacts allow the particular implementation of connectivity at a subcellular level.

## Impact statement

A simple equation linking neurite densities of overlapping neurons to their putative anatomical contacts suggests a potential all-to-all connectivity, typically including sufficient wiring to specifically target individual dendritic compartments.

## Introduction

The functionality of the brain depends fundamentally on the connectivity of its neurons for everything from the propagation of afferent signals (Matthews & Fuchs, 2010; Oh et al, 2014) to computation and memory retention (Hebb, 1949; Hopfield, 1984; Abbott & Regehr, 2004). Connectivity arises from the apposition of complex branched axonal and dendritic arbors which each display a diverse array of forms, both within and between neuronal classes (Bok, 1936; Sholl, 1953; Ascoli et al, 2007, 2008). Despite this complexity, neurons of different classes have been observed to form synapses in highly specific ways, leading to potentially highly structured connectivity motifs within neuronal networks (Binzegger, 2004; Yoshimura & Callaway, 2005; Ohki & Reid, 2007; Perin et al, 2011; Potjans & Diesmann, 2014; Jiang et al, 2015).

Whilst the large-scale EM studies necessary to definitively constrain synaptic connectivity remain prohibitively slow (Briggman & Denk, 2006; da Costa & Martin, 2013; Helmstaedter, 2013) and viral synaptic tracing is limited to small numbers of neurons (Wall et al, 2013), putative synaptic locations from the close juxtaposition of dendrite and axon are more readily measured (Markram et al, 1997; Lee et al, 2016) and provide the potential set of all possible synaptic contacts; the backbone upon which neuronal activity can fine tune connectivity. It has been shown that much of the specificity in putative connectivity can be explained by a detailed analysis of the statistical overlap of different axonal and dendritic arbors (Hill et al, 2012; Markram et al, 2015; Reimann et al, 2017). However such analyses rely on full neuronal reconstructions with large numbers of parameters and are difficult to apply intuitively to microcircuits; there is value in a simple and easily interpretable description of the expected connectivity between a given pair of cells.

The fundamental assumption here is a form of Peters’ Rule, where synapses form uniformly where possible (Peters & Feldman, 1976; Braitenberg & Schüz, 1998). Peters’ rule has been interpreted in a number of different ways at a number of different scales; from predicting of the connectivity between different neuronal classes from their relative abundance (Li et al, 2007) to estimating the number of synapses between a given pair of neurons (Packer et al, 2013). There is experimental evidence both for and against the assumption of uniform synapse formation in different brain regions, species, and under different experimental protocols. A recent review by Rees et al (2017) summarises the experimental evidence for (Packer et al, 2013; van Pelt & van Ooyen, 2013; Merchán-Pérez et al, 2014; Rieubland et al, 2014) and against (Mishchenko et al, 2010; Potjans & Diesmann, 2014; Kasthuri et al, 2015; Lee et al, 2016) Peters’ Rule at the level of individual neurites; but in general it seems that uniform potential structural connectivity is an accurate and powerful model for large regions of the central nervous system.

Given a backbone of neurite structure, neuronal activity is able to strengthen or weaken synapses; allowing memory formation (Hebb, 1949), changes in information storage capacity (Stepanyants et al, 2002; Chklovskii et al, 2004), and sensory tuning (Lee et al, 2016). This relies on the relative dynamism of spine growth and retraction, which occurs on timescales of minutes (Lendvai et al, 2000) (although actual synapse formation can be slower (Knott et al, 2002)), compared to neurite remodelling, which is typically stable over timescales of weeks or months in mature cells (Trachtenberg et al, 2002; Chow et al, 2009). The proportion of close appositions that appear to be bridged by spines at a given time is traditionally referred to as the filling fraction (Stepanyants et al, 2002) and original estimates ranged from 0.1 (macaque V1 visual cortex) to just under 0.4 (rat CA3 hippocampus). A further detail comes from the relationship between the number of anatomical contacts seen under light microscopy with those with synaptic structure under an electron microscope; Markram et al (1997) investigated thick-tufted layer 5 pyramidal cells in rat somatosensory cortex and found that when an axon passed close to a dendrite and displayed substantial swelling indicative of a bouton, then around 80% were true synaptic contacts. Such a reliable and consistent, but not perfect, relationship has since been observed in different neuronal systems (Feldmeyer et al, 1999, 2002; Mishchenko et al, 2010). More recent studies have found, as we observe here in the adult mouse visual cortex dataset of Jiang et al (2015), lower proportions of functional to potential contacts (Kasthuri et al, 2015; Lee et al, 2016). The excess of putative connectivity arises from the extensive branching of cortical neurites and raises questions about the additional functionality provided by this additional metabolic expenditure.

On the postsynaptic side, dendritic trees act as a filter on inputs. The passive electrotonic properties of dendritic cables cause synaptic currents to decay in time and space (Rall, 1964), whilst active processes (Llinas, 1988; Schiller et al, 2000) act to amplify integrated signals that locally exceed a threshold. Inputs to specific regions of some cells allow nonlinear computations to be performed at an intraneuron scale (Mel, 1993; Poirazi et al, 2003; Polsky et al, 2004; London & Häusser, 2005; Losonczy & Magee, 2006). The interaction of clustered or distributed excitatory (Behabadi et al, 2012) and inhibitory (Gidon & Segev, 2012) inputs within a dendritic tree mean that a subcellular-resolution connectome is relevant to realistic network function.

We investigate how well the number of putative synaptic contacts between pairs of neurons can be predicted from simple and intuitive properties of the spatial overlap of their neurite arbours. We find an excess of potential connectivity, both predicted and measured, beyond that described by Stepanyants et al (2002) and sufficient to reliably implement all possible connections at the level of singe cells. We investigate further how well this allows for specific connectivity at the subcellular level of dendritic compartments.

## Results

### Putative synapse number depends on four parameters

A putative synaptic contact is defined as a location where the distance between axon and dendrite is small enough for the gap to be bridged by a dendritic spine. The number of putative synaptic contacts *N* can be estimated by the equation

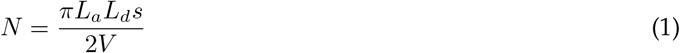

where *L_a_* is the length of axon and *L_d_* the length of dendrite within the axo-dendritic overlap, which has volume *V* . *s* is the maximum spine length at which a synaptic contact could form and typically lies in the range of 1 to 4*μ*m. The full derivation is given in the Methods, but relies on the assumption that straight segments of neurite are distributed at uniform random angles within the overlapping volume and can potentially form a contact if the axon intersects a cylinder of radius *s* around the dendrite (Fig 1a). The form of the equation predicts that the expected number of putative synapses will increase linearly with the maximal spine distance and the lengths of neurite within the axo-dendritic overlap; increasing each of these increases the chance of an axon passing within a maximum spine distance of a dendrite. If these properties are held constant, the expected number of potential contacts decreases with an increase in the volume of the overlapping region due to the reduced density of neurite.

**Fig 1.**
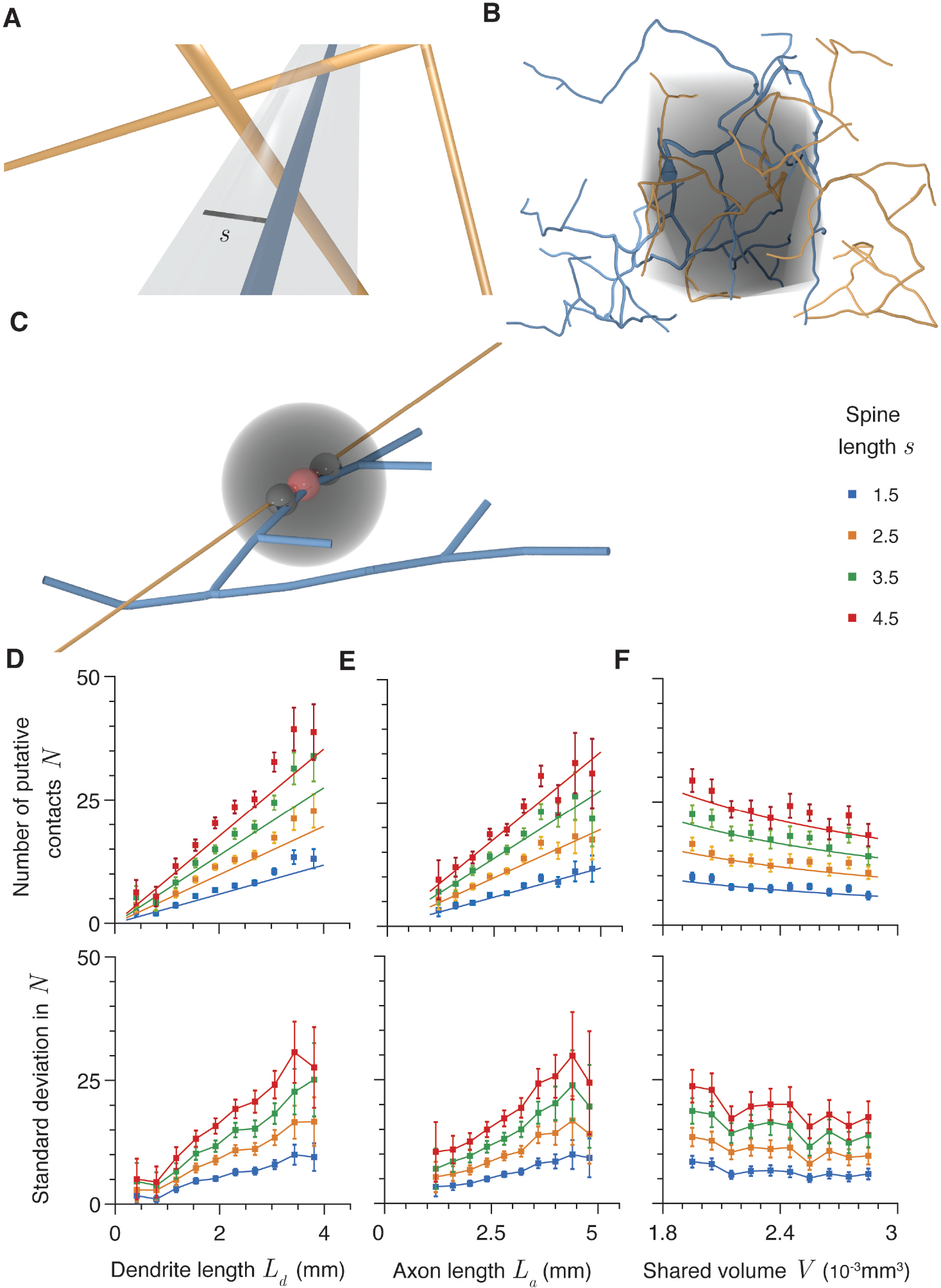
Four factors predict putative synapse number. **A** Schematic illustration of a putative contact between axonal (orange) and dendritic (blue) segments. The grey region is the cylinder of radius *s* (black line) within which putative contacts can form. **B** Shared volume of an example axon (orange) and dendrite (blue). The axo-dendritic overlap is shown in grey. **C** Schematic of region excluded from synapse formation (grey) caused by formation of a potential contact (red). The two black putative contacts are excluded. **D** Expected putative contact number as a function of *L_d_*, dendritic length within the axo-dendritic length. *L_a_* = 3mm and *V* = 2.4 × 10^−3^mm^3^. **E** Expected putative contact number as a function of axonal length within the axo-dendritic length. *L_d_* = 2.4mm and *V* = 2.4 × 10^−3^mm^3^. **F** Expected putative contact number as a function of the volume of the axo-dendritic overlap. *L_d_* = 2.4mm and *L_a_* = 3mm. Different colours show different maximum spine lengths *s*. Error bars show standard error.

The size and shape of the axo-dendritic overlap is therefore a key component of this equation and is defined as follows. Each neuronal arbor is assigned a spanning field, a connected boundary that encompasses all neuronal branches with a tightness dependent on the underlying arbor shape (see Methods and Bird & Cuntz (submitted)). The part of the axonal tree that lies within the spanning field of the dendritic arbor and the part of the dendritic tree that lies within the spanning field of the axonal arbor are used to define the axo-dendritic overlap. A boundary is created around the union of these two sections of arbor with a tightness taken as the mean of those of the two full original trees. This boundary is illustrated by the grey region in Figure 1b and the volume contained within it is defined as *V*.

We apply our equation to estimate the number of putative contacts between generalised minimum spanning trees Cuntz et al (2010) that reproduce the properties of axonal and dendritic trees, where the measured number is denoted *n*. In order to prevent an unbounded clustering of synaptic contacts whenever an axon and dendrite pass close together at a single point, we further introduce an exclusion region around each contact (illustrated by the grey sphere in Fig 1c). The closest apposition between dendrite and axon is selected as a contact and all other appositions within a certain distance, typically 3*μ*m, are excluded from forming putative contacts (illustrated by the small black spheres in Fig 1c). The closest remaining apposition is then selected and another exclusion applied. This is repeated until there are no appositions closer than *s* remaining. The exclusion region is a conservative constraint as Schmidt et al (2017) found that around 20% of synaptic connections in rat medial entorhinal cortex exhibited clustering, with mean intercontact distances within a cluster of 3.7 *μ*m and 4.8 *μ*m onto excitatory and inhibitory dendrites respectively. The number of synaptic contacts given by this algorithm is therefore likely to be an underestimate of the true number, but ensures that *n* does not depend strongly on the sampling frequency of the neurite discretisation (or grow to infinity if the neurites are treated continuously). In terms of postsynaptic functionality, tightly clustered contacts are far more likely to innervate a single dendritic compartment and so provide a strong but spatially localised input that does not alter the connectivity structure at the subcellular level.

Figures 1d to f plot the number of putative synaptic contacts found numerically for synthetic neuronal arbors generated using generalised minimum spanning trees for different maximum spine lengths *s* as a function of each of *L_d_*, *L_a_*, and *V* when the other parameters are held approximately constant. The dashed lines give the predictions of Eq 1 in each case and show a good match between theory and simulation for these synthetic neurites. The standard deviations are plotted below the mean in each case and are quite large, growing proportionally with the mean.

Eq 1 is similar to results introduced by Stepanyants et al (2002) for synaptic contacts onto a given dendritic tree by all axons in a tissue and Chklovskii (2004) to determine the total number of afferent synapses onto a particular dendrite given the total abundance of axons within a cortical column. The application here differs from previous usage as it explicitly accounts for an individual axonal tree, allowing for estimation of cell-type specific connectivity given the statistics of axo-dendritic pairs. It is also a simplification of the detailed approaches to estimating pairwise connectivity in Hill et al (2012), Markram et al (2015), and Reimann et al (2017) as well as the dendritic-density based approaches of Liley & Wright (1994), Amirikian (2005), van Pelt & van Ooyen (2013), and Aćimović et al (2015). By accurately modelling potential connectivity in terms of four simple parameters, this equation simply and robustly highlights the major determinants of microcircuit structure.

### Other factors do not substantially influence putative synapse number

Neurites take a very wide array of shapes with different branching statistics and locations relative to one another (Bok, 1936; Ascoli et al, 2007, 2008); although we have shown a relationship between four features of a neuronal pairing and the expected putative synapse number *N*, it is worthwhile to consider whether other factors implicit to our model may have an effect. We have therefore investigated whether three other features alter the accuracy of the prediction under Eq 1. Firstly we considered whether the shape of the domain used to create the synthetic neurites has an impact. For Figure 1, the generalised minimum spanning tree algorithms were used to generate trees within cubes, but various other shapes such as cones, spheres, and cylinders do not affect the results (Fig 2a).

**Fig 2.**
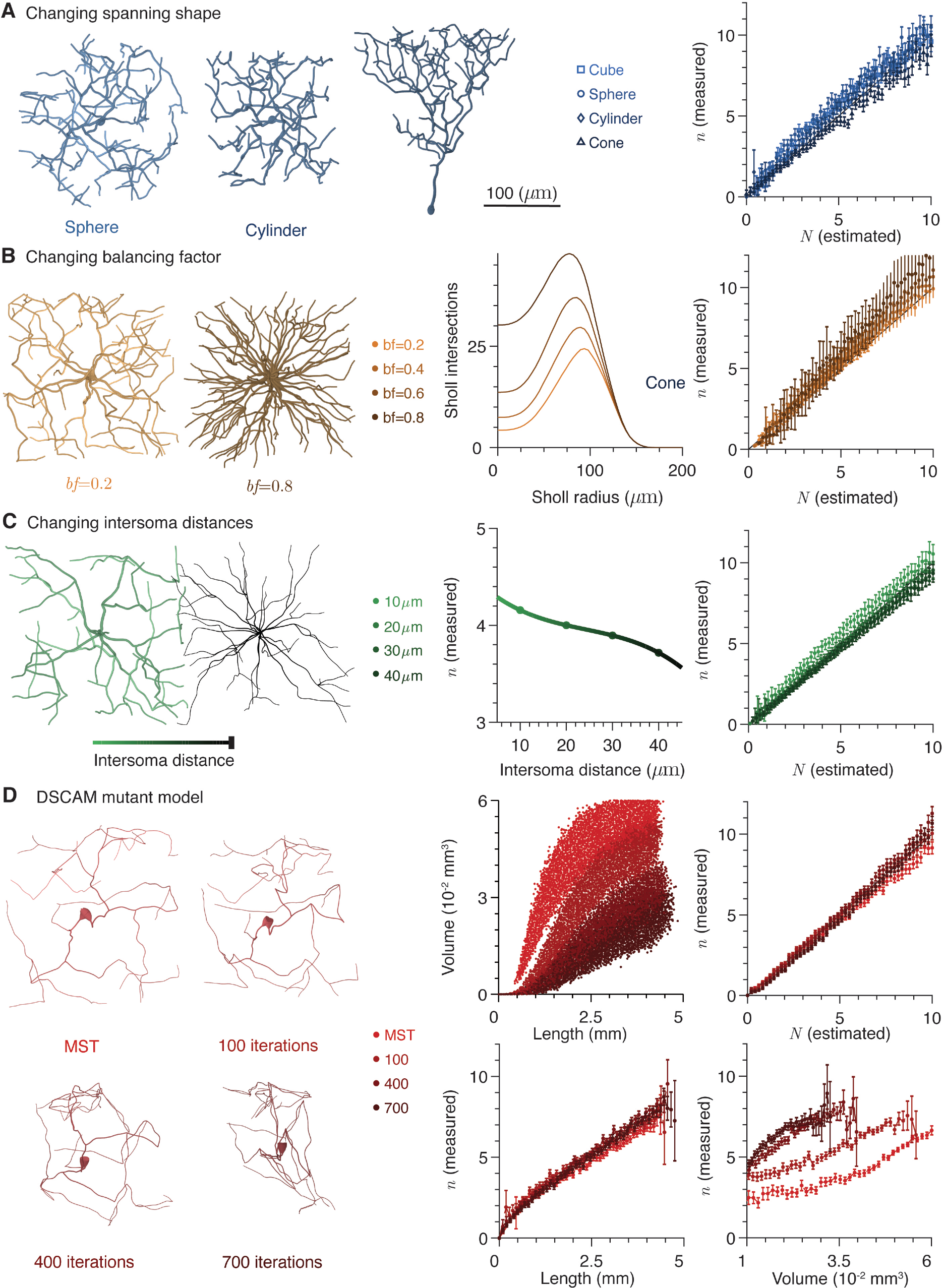
Other factors do not influence putative synapse number (continued) **A** Left: Example morphologies of neurites grown in different domains; from left to right, sphere, cylinder, and cone. Right: Measured putative contact number as a function of estimated putative contact number for different neurite domains. **B** Left: Example morphologies generated with different balancing factors; left *bf* = 0.2 and right *bf* = 0.8. Scale bar as above. Centre: Mean Sholl intersection profiles for neurons with different balancing factors. Right: Measured putative contact number as a function of estimated putative contact number for different balancing factors. **C** Left: Schematic of intersoma distance. Centre: Mean expected numbers of contacts as a function of intersoma distance. Right: Measured putative contact number as a function of estimated putative contact number for different intersoma distances. **D** Left: Example morphologies with different numbers of iterations of the DSCAM null algorithm: 0 (generalised MST), 100, 400, and 700. Centre top: Volume spanned by a dendrite as a function of length for different numbers of iterations of the DSCAM null algorithm. Right top: Measured putative contact number as a function of estimated putative contact number for different numbers of iterations of the DSCAM null algorithm. Centre bottom: Expected number of putative contacts as a function of dendrite spanning volume. Right bottom: Expected number of putative contacts as a function of dendrite length. In all cases, error bars show standard error.

The balancing factor *bf* in the generalised minimum spanning tree model determines the balance between costs associated with additional neurite length and conduction delays caused by long path distances between synapses and the soma. A balancing factor of zero corresponds to a pure MST where conduction delays are ignored and in the limit of high balancing factors, all synapses are directly connected to the soma. Typically real non-planar neurons have dendrites with balancing factors in the range 0.2 to 0.8 (Cuntz et al, 2010) and there is a roughly exponential relationship between increasing balancing factor and the centripetal bias as quantified by the root angle distribution (Bird & Cuntz, submitted). We typically set the balancing factors of the dendrite and axon to 0.2 and 0.7 respectively to account for the different features of these neurites (Cuntz et al, 2007; Budd et al, 2010; Teeter & Stevens, 2011). Varying the dendritic balancing factor over the range 0.2 to 0.8, the majority of the range observed in reconstructed neurons, whilst keeping other features the same does not alter the accuracy of the predictions of Eq 1 (Fig 2b). This result is particularly surprising as we have recently shown that different balancing factors lead to substantially different distributions of neurite mass within their spanning fields (Fig 2b, centre) and violates the assumption of isotropically distributed branches through its effect on the root angle distribution.

Finally, the inter-soma distance does not matter as long as the cable lengths and overlapping volume are controlled for (Fig 2c). This last point is particularly interesting, as a number of studies report a strong influence of intersoma distance on predicted connectivity (Hellwig, 2000; Kalisman et al, 2003; van Pelt & van Ooyen, 2013); we find that intersoma distance only matters through the negative correlation between distance and the neurite lengths within an overlapping volume.

### Clustered dendrites modelling the DSCAM-null mutation do not cause a loss of potential connectivity

The above results are for dendrties with the properties of relative space-filling and spatial uniformity particular to trees that minimise metabolic costs (Cuntz et al, 2007; Wen et al, 2009; Bird & Cuntz, submitted). A major exception to these properties comes from invertebrate neurons with a Down Syndrome Cell Adhesion Molecule (DSCAM) null mutation (Schmucker et al, 2000). The inactivation of this gene reduces the self-avoidant tendency of neurites and leads to pathologically clustered dendrites (Soba et al, 2007). In Methods we describe a modification of the existing MST model to produce artificial neurites that have the characteristics of DSCAM null mutants. In short, the algorithm iteratively randomly selects branches of the neurite and moves them 10% closer to the closest neighbouring branch. Iterating this process produces progressively more clustered morphologies (Fig 2d, left).

Increasing the number of iterations changes the relationship between length and volume as dendrites become more densely clustered (Fig 2d, centre top). However, when applied to such arbors, the predictions of Eq 1 still hold (Fig 2d, right top). This means that the potential connectivity of neurites that do not effectively fill space remains predictable from Eq 1 and that pathological clustering of dendrites does not cause loss of function through lost connectivity beyond that predicted by changes in dendrite length and spanning field (Mychasiuk et al, 2012). This is a slightly counterintuitive finding as Wen et al (2009) found that non-pathological dendritic branching statistics are in line with those that maximise the connectivity repertoire of afferent connections. The null mutation causes a greater density of dendrite within its spanning field and so DSCAM mutants have a relatively high number of putative contacts within a given spanning volume (Fig 2d, centre bottom). However, this is balanced by the reduced amount of axon that typically intersects the dendritic spanning field and so the null mutation has no effect on the relationship between the length of the dendritic tree and the expected number of contacts it receives (Fig 2d, right bottom).

### Synapse estimation for reconstructed morphologies

Our model produces accurate estimates of putative synaptic contact number and connection probability for the generalised MST models that accurately simulate real neurites, but also applies directly to neural reconstructions. To demonstrate this, we consider a dataset of reconstructions from the rat barrel cortex and developmental subplate by Marx et al (2017). When neurons are randomly paired with random offsets in their somata and orientation (see Methods and Fig 3a), Eq 1 correctly predicts the number of putative axo-dendritic synaptic contacts (Fig 3b). The distribution of measured values of *n* for each expected value *N* are shown by the heatmap in Figure 3c.

**Fig 3.**
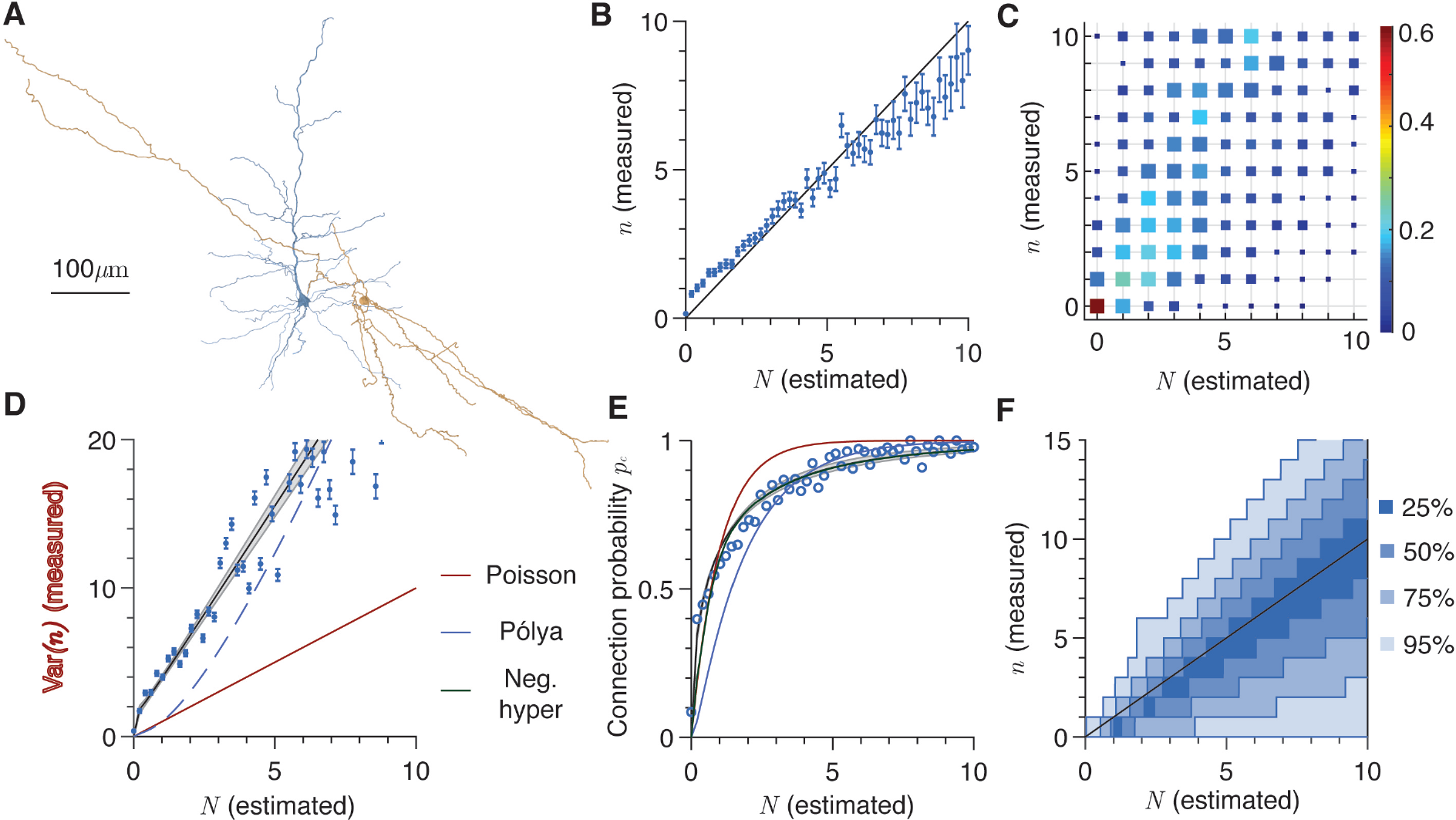
Predictions for reconstructed morphologies. **A** Example of dendritic (blue) and axonal (orange) morphologies at an arbitrary displacement and orientation (Ascoli et al, 2007; Marx et al, 2017). **B** Mean measured versus estimated putative contact number (blue markers and error bars). Equality is given by the black line. Error bars show standard error. **C** Probability distribution of putative contact numbers for each integer interval of estimated putative contact number. Distributions are normalised for each estimated interval and square sizes scale linearly with the occurrence of each probability in the grid. **D** Variance in measured versus estimated mean putative contact number (blue markers and error bars). The Poisson model variance is shown by the solid red line and the best fit by the solid black line with the 95% confidence interval in grey (coefficients are 2.937 (2.8, 3.146)). The dashed blue line shows the Pólya model with parameters fitted to the connection probability (Eq 3). Error bars show standard error. **E** Connection probability as a function of estimated putative contact number. The fits from the Poisson, Pólya, and negative hypergeometric models are shown by the red, blue, and green lines respectively. The fit from Eq 3 is shown by the solid black line and grey shaded region. **F** Confidence intervals (25%, 50%, 75%, and 95%) for values of *n* as a function of *N* under the negative hypergeometric model (Eq 16). In all panels, maximum spine distance *s* = 2.5*μ*m.

It is interesting to note the variability in these results. Figures 1d to f shows the variance in the measured value of *N* as a function of the underlying parameters and it typically takes large values. Similarly, Figure 3d shows the distribution of measured values of *n* for each estimated integer value of *N* . Both illustrate the large variation in possible true numbers of putative contacts for a given set of parameters *L_a_*, *L_d_*, and *V* . The wide variability here means that Eq 1 is unlikely to be perfectly accurate when applied to a single axo-dendritic pair, but is a true estimate of the expected number of putative synapses using relatively simple parameters.

### Connection probability *p_c_* and the distribution of *n*

In addition to the expected number of potential connections, the connection probability *p_c_* (the probability that *n* > 0), is important for inference of network structure. If the putative contacts formed independently with a fixed probability, then the probability distribution of measured anatomical contacts for a given value of *N* under Eq 1 would take a Poisson distribution (Eq 11) with mean and variance both given by *N* . However, the variance in the measured numbers of putative contacts typically exceeds the mean (Fig 3d) and so makes the assumption of contacts forming independently untenable. van Pelt & van Ooyen (2013) found a similar effect when estimating connection probability from their density based model; correlations in putative contact formation arise from the fact that both neurites are connected trees and so close appositions in one location can increase the chance of more close appositions occurring. Indeed, the connection probability given by the Poisson model fitted to the mean

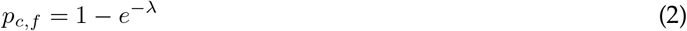

where *λ* = *N*, does not match the measured probabilities (Fig 3e, blue line). The measured connection probability *p*_*c,measured*_ is better described by an equation of the form

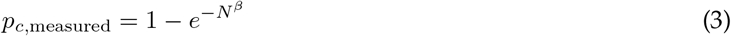

where the fitted value of *β* is 0.5437, with a 95% confidence interval of (0.5069, 0.5805) (Fig 3E, black line and shaded area).

The Pólya distribution (Eq 12) modifies the Poisson distribution to allow the mean and variance to differ and can be used to describe correlated occurrences (Blom et al, 1993). The variance in *n* grows faster than *N* and is well-described by a function of the form var(*n*) = *aN* + *N^b^* where the parameters are (with 95% confidence intervals) *a* = 2.944 (2.769, 3.119) and *b* = −0.124 (−0.246, −0.001). This is plotted as the solid black line and shaded grey area in Figure 3d. The second term *N^b^* is necessary to capture the initial growth in the variance for small values of *N* that is particularly apparent when plotting the Fano factor var(*n*)/*N* in Figure S3b. It should be noted that the variance in *n* as a function of *N* is fundamentally different to the variances in *n* as functions of *L_a_*, *L_d_*, and *V* shown in Figure 1. The estimates of each value of *N* come from a wide variety of possible combinations of the underlying parameters that obey Eq 1; the resultant variance in *n* therefore has a complex dependence on the underlying factors, weighted by their joint likelihood of occurrence, that is best described empirically.

Fitting the Pólya distribution to the mean and variance gives the connection probability as

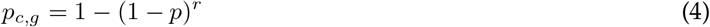

where the parameters are given by *p* = 1 − 1/(*a* + *N*^*b*−1^) and *r* = *N*/(*a* − 1 + *N*^*b*−1^). However, although this gives a better estimate of *p_c_* than Eq 2 for large values of *N*, it is even less accurate for *N* ⪅ 2 (Fig 3e, blue line). This is because the increased variance moves probability mass away from the mean value approximately symmetrically and so increases the mass at 0 in contrast to the measurements (Fig 3c). It is also possible to fit the connection probability of the Pólya distribution to *p*_*c,measured*_ exactly (Eq 13), but this leads to an inaccurate estimate of the variance (Fig 3d, blue dashed line).

To better capture the measured properties of the distribution of *n* given an estimate *N*, we therefore use a three-parameter negative hypergeometric distribution (Eq 16) to describe the distribution of *n* for small values of *N* (less than 10). The negative hypergeometric distribution can be more closely matched to the moments and connection probability of the observed distributions and in particular has connection probability in terms of its parameters Δ, *K*, and *ρ*

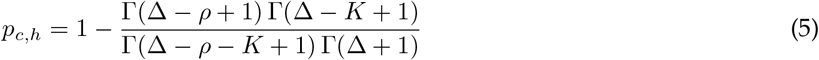

where 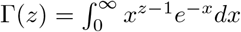 is the gamma function. For larger values of *N* (greater than 10), the Pólya model, fitted to the mean and variance is a good description of the data (Fig S3). The Pólya distribution typically describes the probability of a number of events occurring, when each occurrence increases the likelihood of subsequent events. This is an appropriate model for *n* as the spatial correlations within connected neurites mean that a single close apposition increases the chance of neighbouring regions of axon and dendrite also lying close together. The negative hypergeometric distribution can be interpreted as a generalisation of the Pólya distribution to the case where the total number of possible occurrences is limited. In the neurite case this means that a close apposition can increase the probability of more close appositions locally, while globally reducing the probability of more close apposition as it accounts for some proportion of the total available cable. This is particularly important for smaller values of *N*, when *L_a_* and *L_d_* are likely to be relatively small.

### Synapse estimation for reconstructed microcircuits

In the previous sections, we considered the number of putative contacts between reconstructed morphologies with random somatic locations. This verified the predictions of the generalised minimum spanning tree model for real axons and dendrites, but left open the question of how the specific arrangement of axonal and dendritic spanning fields within cortical circuits can lead to certain connectivity patterns. Jiang et al (2015) produced a dataset of reconstructions from the visual cortex of adult mice, and in particular often reconstructed multiple cells from the same slice (Fig 4a), allowing Eq 1 to be tested on a large set of cells in context with one another. The predictions hold very well, allowing accurate predictions of both overall (Fig 4b) and cell-type specific (Fig 4c) putative connectivity.

**Fig 4.**
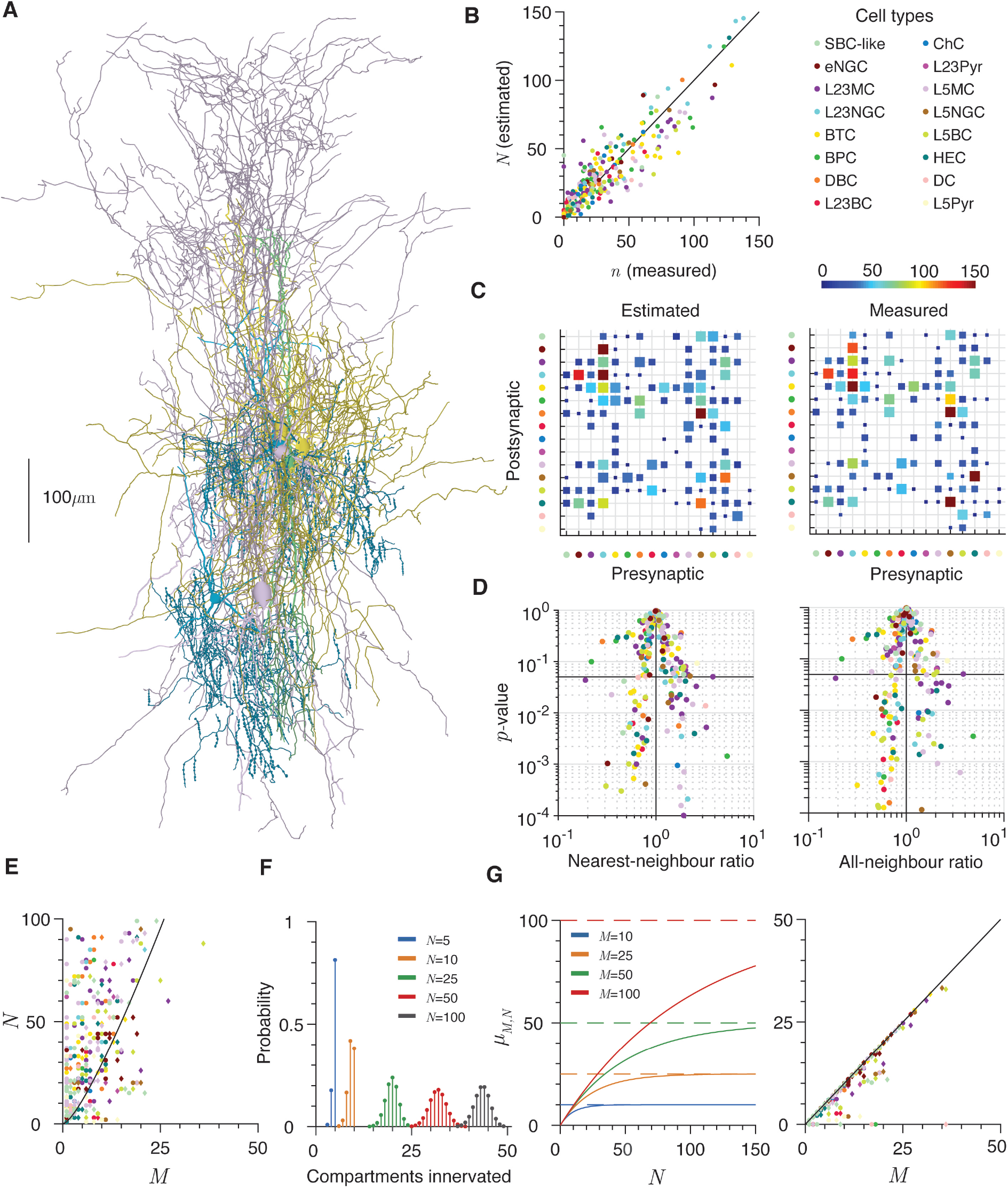
Excess putative connections in a reconstructed microcircuit. **A** Example of 7 cell types reconstructed from the same slice (Ascoli et al, 2007; Jiang et al, 2015). Cells are, using the definitions in Jiang et al (2015), L2/3 bitufted cell (2 examples), L2 Martinotti (2 examples), L2/3 chandelier (2 examples), and L2/3 bipolar (1 example). Diameters are increased by 1*μ*m to increase visibility and morphologies are coloured by cell class (see below). **B** Measured putative connectivity as a function of estimated putative connectivity for the microcircuit data. Colours correspond to postsynaptic cell type (see legend). Maximum spine distance *s* = 3*μ*m. **C** Predicted (left) and measured (right) cell-type specific connectivity. The horizontal axis shows the pre- and the vertical axis the post-synaptic cell types and contact numbers are per connected pair. Square sizes scale linearly with the occurrence of each connectivity in the grid. Maximum spine distance *s* = 3*μ*m. **D** Nearest-(left) and all-(right) neighbour ratios plotted against the two-sided *p*-value for each cell pair. The horizontal lines show *p* = 0.05 and the vertical lines a ratio of 1. **E** Number of afferent contacts *N* as a function of the number of dendritic compartments *M* that could be innervated in the axo-dendritic overlap. Colours indicate dendrite type; circles correspond to the branch estimate of *M* and diamonds to the electrotonic estimate. The black line shows the expected number of contacts necessary to innervate every compartment (Eq 4). **F** Examples of the distributions of distinct compartments (out of 50) innervated by *N* = 5, 10, 25, 50, and 100 contacts (Eq 5). **G** Left: Expected number *μ_n,M_* of distinct compartments innervated as a function of the number of contacts *N* for different numbers of compartments *M* = 10, 25, 50, and 100 (Eq 6). Solid lines show *μ_n,M_* and dashed lines show *M*, the maximum possible number in each case. Right: Expected number of innervated compartments *μ_n,M_* as a function of the number of dendritic compartments *M* that could be innervated in the axo-dendritic overlap (Eq 6). Colours indicate dendrite type; circles correspond to the branch estimate of *M* and diamonds to the electrotonic estimate. The black line shows *M*, the maximum number of compartments that could be innervated in each case.

### Putative contact numbers often exceed those necessary for reliable connectivity

The numbers of putative contacts from both Eq 1 and direct measurements are often very high in this dataset. Experimental studies find many fewer functional contacts, often an order of magnitude lower, between cell pairs (Markram et al, 1997; Kasthuri et al, 2015; Lee et al, 2016). An initial hypothesis would be that very high numbers of potential contacts are necessary to increase the probability of having at least one potential connection in order to allow a microcircuit to function. Under the model of Eq 3, we estimate that the probability that the cell pair with the greatest number of putative synapses, an elongated (L1) neurogliaform to L2/3 neurogliaform cell, would be disconnected given their neurite lengths within the overlapping region is or less than one in 3 million.

If the axon length within the overlapping volume were to half, the probability of no connection would still be one in thirty thousand. For the tenth most putatively connected cell pair, a pair of L2/3 double bouquet cells, the probabilities decreases from one in two hundred thousand to one in five thousand. These probabilities are not that low given the number of cells within a cortical column, but do suggest that the metabolic cost to reliably establish single connections is far below that typically paid by these cells. It should also be noted that slicing artefacts, by removing neurite outside of the slice, will tend to bias the numbers of putative contacts recorded here down (Jiang et al, 2016); in intact cortex the putative connectivity will be at least as high as that observed here. Our findings are in line with the very high degree of synaptic redundancy observed by Kasthuri et al (2015) in mature mouse somatosensory cortex and Lee et al (2016) in mature mouse visual cortex.

### Putative contacts are well-distributed within the axo-dendritic overlap

To determine the distribution of putative connections within the axo-dendritic overlap, and in particular whether they are more clustered or more regular than a uniform random spatial distribution (ie a homogeneous spatial Poisson process, see Methods), we used both the nearest-neighbour ratio (NNR) and the all-neighbour ratio (ANR) (Chandrashekhar, 1943). The nearest-neighbour ratio quantifies whether the distance between a potential contact and the closest other contact is more or less than would be expected for a spatially homogeneous random process. A nearest-neighbour ratio of one implies that the potential contacts are distributed within the axo-dendritic overlap precisely as one would expect from a homogenous Poisson process, whereas a ratio of less than one implies clustered and more than one well-distributed potential contacts. The number of potential contacts varies widely between cell pairs, so *p*-values (see Methods) are plotted against the nearest-neighbour ratio in Figure 4d to indicate the significance of the difference from one for each cell pair. Colours in this panel indicate the cell type of the presynaptic neuron.

As synaptic contacts are distributed along neurite arbors, there is potential for local spatial correlations to arise and dominate the pairwise measure given by the nearest-neighbour ratio. To determine whether the contacts display local correlation along arbors, but more general independence, we also computed the all-neighbour ratio: the average deviation of each putative contact from the centroid of all contacts. This measure is less sensitive to local correlation and is plotted in Figure 4d.

Both measures show that there is a spread of ratios; 35 of 233 pairs with more than one putative contact are significantly clustered and 31 are significantly regular under the nearest-neighbour ratio, with numbers of 53 and 35 for the all-neighbour ratio. However there appears to be no significant and consistent clustering or regularity by either pre- or postsynaptic cell types (see Table 2 for mean nearest-neighbor values and *p*-values by presynaptic class). This lack of apparent spatial structure in putative connections between a cell pair is an interesting intermediate case. Merchán-Pérez et al (2014) found uniform randomness in the location of all synapses within a volume of neuropil without reference to specific neurites, while synapses along a given axon or dendrite will be clearly spatially correlated. van Pelt & van Ooyen (2013) concluded that spatial correlations were the cause of the mismatch between their model of density-based mean putative contact number predicition and the connection probability and we have observed a similar effect (Fig 3e). However, they did not directly check for spatial correlations in putative contacts and appear to use lower neurite densities than those seen in the reconstructed data here, which could enhance the impact of potential contacts sharing a branch. Of particular interest here is that inhibitory cell types do not appear to form potential contacts in a more spatially structured way than excitatory cells (see Table 2). This is despite the fact that cortical inhibitory neurons stereotypically innervate specific regions of excitatory cells (Ascoli et al, 2008; Hill et al, 2012). These results suggest that such specificity could come entirely from the axonal growth region rather than individual local targeting processes.

**Table 1.** Table summarising symbols and terms.

| Symbol | Interpretation |
| --- | --- |
| $L_a$ | Axon length within overlapping region |
| $L_d$ | Dendrite length within overlapping region |
| $M$ | Number of dendritic compartments |
| $n$ | Measured number of putative contacts |
| $N$ | Estimated number of putative contacts |
| $N_{\text{Complete}}$ | Number of putative synaptic contacts to innervate all dendritic compartments |
| $p_c$ | Probability that a pair of cells are connected (second subscript denotes distribution model) |
| $s$ | Maximum spine length |
| $V$ | Volume of axo-dendritic overlap |
| $\mu_{M,n}$ | Expected number of dendritic compartments (out of $M$) innervated by $n$ synaptic contacts |

**Table 2.** Table of in-context reconstructions of the dataset from Jiang et al (2015). Columns from left to right are: Cell type, number of individual morphologies, total number of cell pairs involving neurons of this class, mean number of efferent synapses, mean nearest-neighbour ratio (NNR) of efferent synapses, mean *p*-value significance that this ratio is different from one, and mean number of afferent synapses (with 95% confidence interval). ± shows the standard error over different cells of each class and confidence intervals are for the mean of each class using Eqs 12 and 16. Columns from left to right are: Cell type, mean total number of dendritic compartments *M*, mean number of dendritic compartments *M* that lie within the axo-dendritic overlap, ratio of number of afferent synapses *n* to the number necessary to expect to innervate each available dendritic compartment *N*_Complete_ (with 95% confidence interval), and ratio of mean number of available compartments innervated *μ_n,M_* to number of available of compartments *M* (with 95% confidence interval). For the last four columns, the upper values come from the branch-based estimate of compartments, and the lower from the attenuation-based estimate. ± shows the standard error over different cells of each class and confidence intervals are for the mean of each class using Eqs 12 and 16.

| Cell type | Numbers |  | Efferent contacts |  |  | Afferent contacts |
| --- | --- | --- | --- | --- | --- | --- |
| | Examples | Pairs | $n_{\text{efferent}}$ | NNR | $p$-value | $n_{\text{afferent}}$ |
| SBC-like | 21 | 22 | $25.79 \pm 7.66$ | $1.04 \pm 0.18$ | $0.39 \pm 0.12$ | $37.43 \pm 9.05$ (23.39, 53.54) |
| eNGC | 21 | 17 | $58.05 \pm 11.67$ | $0.89 \pm 0.11$ | $0.31 \pm 0.14$ | $65.55 \pm 12.65$ (44.23, 89.55) |
| L23MC | 20 | 51 | $39.88 \pm 4.89$ | $1.36 \pm 0.12$ | $0.29 \pm 0.05$ | $45.01 \pm 4.79$ (27.85, 64.56) |
| L23NGC | 20 | 28 | $67.4 \pm 11.6$ | $1.12 \pm 0.09$ | $0.39 \pm 0.09$ | $92.54 \pm 11.12$ (65.97, 121.71) |
| BTC | 20 | 50 | $54.13 \pm 6.37$ | $0.94 \pm 0.05$ | $0.39 \pm 0.06$ | $40.16 \pm 5.46$ (24.42, 58.27) |
| BPC | 20 | 54 | $43.32 \pm 5.61$ | $1.29 \pm 0.43$ | $0.3 \pm 0.09$ | $23.51 \pm 4.11$ (13.28, 35.3) |
| DBC | 20 | 15 | $64.71 \pm 16.36$ | $1.04 \pm 0.16$ | $0.34 \pm 0.11$ | $63.29 \pm 13.96$ (43.19, 85.86) |
| L23BC | 15 | 16 | $52.79 \pm 13.65$ | $0.82 \pm 0.08$ | $0.24 \pm 0.1$ | $53.17 \pm 13.77$ (36.21, 72.75) |
| ChC | 20 | 15 | $26.24 \pm 8.45$ | $1.09 \pm 0.1$ | $0.47 \pm 0.12$ | $19.47 \pm 5.58$ (8.76, 32.35) |
| L23Pyr | 20 | 9 | $29.38 \pm 9.61$ | $1.36 \pm 0.33$ | $0.61 \pm 0.27$ | $28.46 \pm 9.29$ (15.92, 43.46) |
| L5MC | 21 | 42 | $33.69 \pm 5$ | $1.31 \pm 0.08$ | $0.39 \pm 0.07$ | $39.63 \pm 5.09$ (23.54, 58.27) |
| L5NGC | 21 | 19 | $69.24 \pm 17.55$ | $1.26 \pm 0.15$ | $0.28 \pm 0.13$ | $67.84 \pm 15.58$ (48.6, 89.24) |
| L5BC | 21 | 36 | $50.43 \pm 8.58$ | $0.94 \pm 0.08$ | $0.28 \pm 0.06$ | $54.37 \pm 8.58$ (35.39, 75.96) |
| HEC | 21 | 32 | $54.71 \pm 10.06$ | $1.11 \pm 0.14$ | $0.14 \pm 0.03$ | $56.78 \pm 9.92$ (38.44, 77.78) |
| DC | 21 | 19 | $23.58 \pm 7.87$ | $1.23 \pm 0.34$ | $0.33 \pm 0.13$ | $36 \pm 9.39$ (22.58, 51.58) |
| L5Pyr | 21 | 9 | $42.18 \pm 20.04$ | $2.63 \pm 0$ | $0.09 \pm 0$ | $40.36 \pm 20.16$ (27.36, 55.55) |

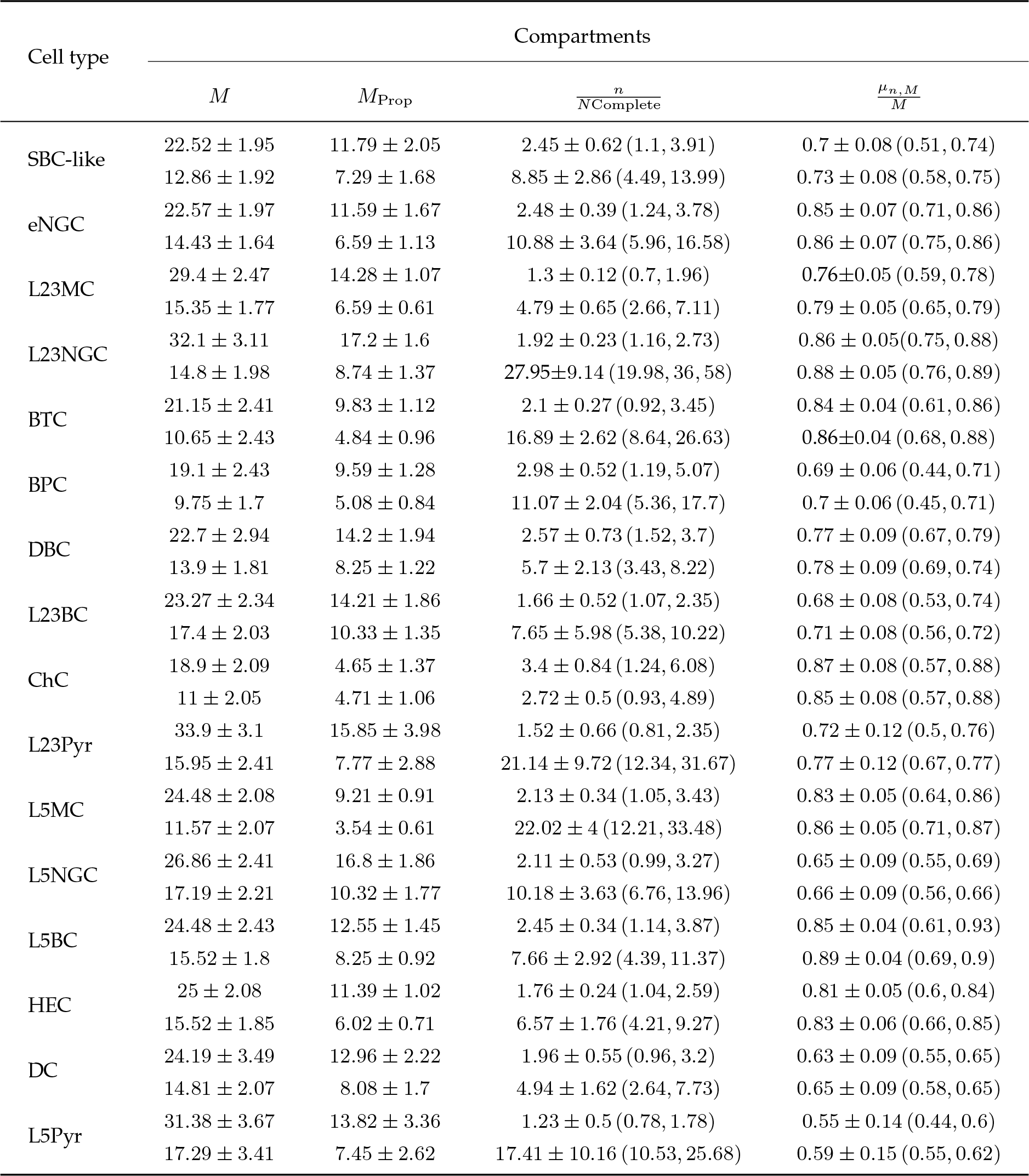

### Excess potential connections allow for the innervation of multiple dendritic compartments

Given that expected potential contact numbers are far in excess of that necessary to produce reliable connectivity at the cellular level, and that contacts lack apparent spatial structure, it is both informative and feasible to investigate how cortical neurite densities can implement sub-cellular connectivity by innervating specific or distinct dendritic compartments. Definitions of dendritic compartments vary in the literature (Mel, 1993; Poirazi et al, 2003; Polsky et al, 2004; London & Häusser, 2005; Branco & Häusser, 2010; Cuntz et al, 2010; Behabadi et al, 2012) and certainly the electronic structure of a dendritic tree changes dynamically with network activity due to both synaptic activation and thresholded processes (Llinas, 1988; Schiller et al, 2000; Gidon & Segev, 2012; Ferrarese et al, 2018). To investigate compartmentalisation, we consider two simple descriptions of a dendritic compartment. The first is simply that each individual dendritic branch is a compartment (Branco & Häusser, 2010) and the second assigns reasonable passive electrotonic properties (see Methods) to a dendrite and divides the tree into regions within which synaptic currents do not attenuate below a certain threshold (Cuntz et al, 2010). The first estimate is indicated by diamonds and the second by circles in Figures 4e and 4g. These are both certainly drastic simplifications of the true integrative properties of a dendrite, particularly within an active microcircuit, but provide an informative first step. To account for the fact that the axo-dendritic overlap only covers a part of the dendrite, we scale the compartment number down to match the proportion of dendrite that lies within this volume.

The spatial spread of potential contacts means that it is reasonable to assume that they randomly innervate dendritic compartments independently and uniformly. In this case the expected number of contacts necessary to contact every one of *M* compartments is given by the standard solution to the coupon collector’s problem (see Methods)

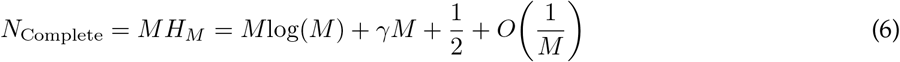

where *H_M_* is the *M*-th harmonic number and *γ* ≈ 0.57721 is the Euler-Mascheroni constant. The second equality relies on the asymptotics of harmonic numbers. Plotting *M* against *n* (Fig 4e, black line corresponds to Eq 6) for the cells in this dataset shows that most neurons receive enough putative contacts to expect to innervate each of the available dendritic compartments. In Table 2, the ratio of putative contacts to *N*_Complete_ is above one on average for every cell class considered, even when each individual branch is treated as a distinct dendritic compartment.

The second quantity to consider is the distribution of the number of dendritic compartments innervated by at least one synapse. The probability mass function for this quantity, given *M* compartments and *n* contacts, is

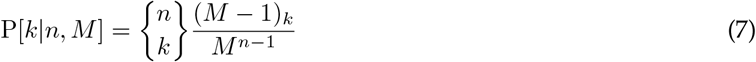

where 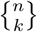 is a Stirling number of the second kind (Abramowitz & Stegun, 1965) and (*M* − 1)*_k_* is the falling factorial (*M* − 1)*_k_* = (*M* − 1)(*M* − 2) · · · (*M* − *k* + 1). This distribution has intuitive properties (Fig 4f): when *n* is much larger than *M*, then probability mass is grouped around *M* as it is highly likely that every compartment will be innervated. Conversely, when *M* is much larger than *n*, the probability mass is grouped around *n* as it is likely that every synapse will innervate a distinct compartment. The mean number *μ_M,n_* of distinct compartments innervated is given by

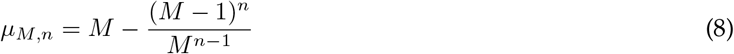

Figure 4g (left side) plots *μ_M,n_* as a function of *n* for different values of *M*, highlighting the asymptotic behaviour as *n* grows larger than *M* . The right-hand side of Figure 4g plots *μ_M,n_* against *M* for the cells in this dataset, given the number of putative contacts they receive. Many cell pairs have sufficient putative contacts to expect to be able to innervate almost all of the available dendritic compartments. In Table 2, the ratio of *μ_M,n_* to *M* is frequently above 0.75 when averaged over each cell class, meaning that three out of four available compartments could potentially receive a synaptic contact.

### Applying the prediction of putative connectivity

As a final demonstration of the utility of Eq 1 in predicting putative connectivity, we show how the neurite lengths and shared volumes, as well as the expected number of putative synapses, vary with intersoma distance for the reconstructed cortical morphologies of mouse (Fig 5a, top) and human (Fig 5a, top) cells published by the Allen Institue for Brain Science (Allen Brain Institute, 2015). These cells are not typically imaged in context, but do have the depth of the soma below the cortical surface reliably recorded, alongside the orientation within the slice (see Fig S5). For the demonstration in Figure 5, we randomly pair axonal and dendritic reconstructions using their recorded cortical depth and a random offset in the plane parallel to the cortical surface (see Methods). We can see a decrease in all three relevant parameters *L_a_*, *L_d_*, and *V* with increasing intersoma distance (Fig 5b, top three panels), and the fact that the expected number of putative contacts depends on the product of *L_a_* and *L_d_* over *V* means that this is accompanied by a decrease in *N* (Fig 5b, bottom panels). These predictions give an intuition of the structural backbone to functional connectivity in cortex that can be progressively refined as more reconstructions of individual cell types appear.

**Fig 5.**
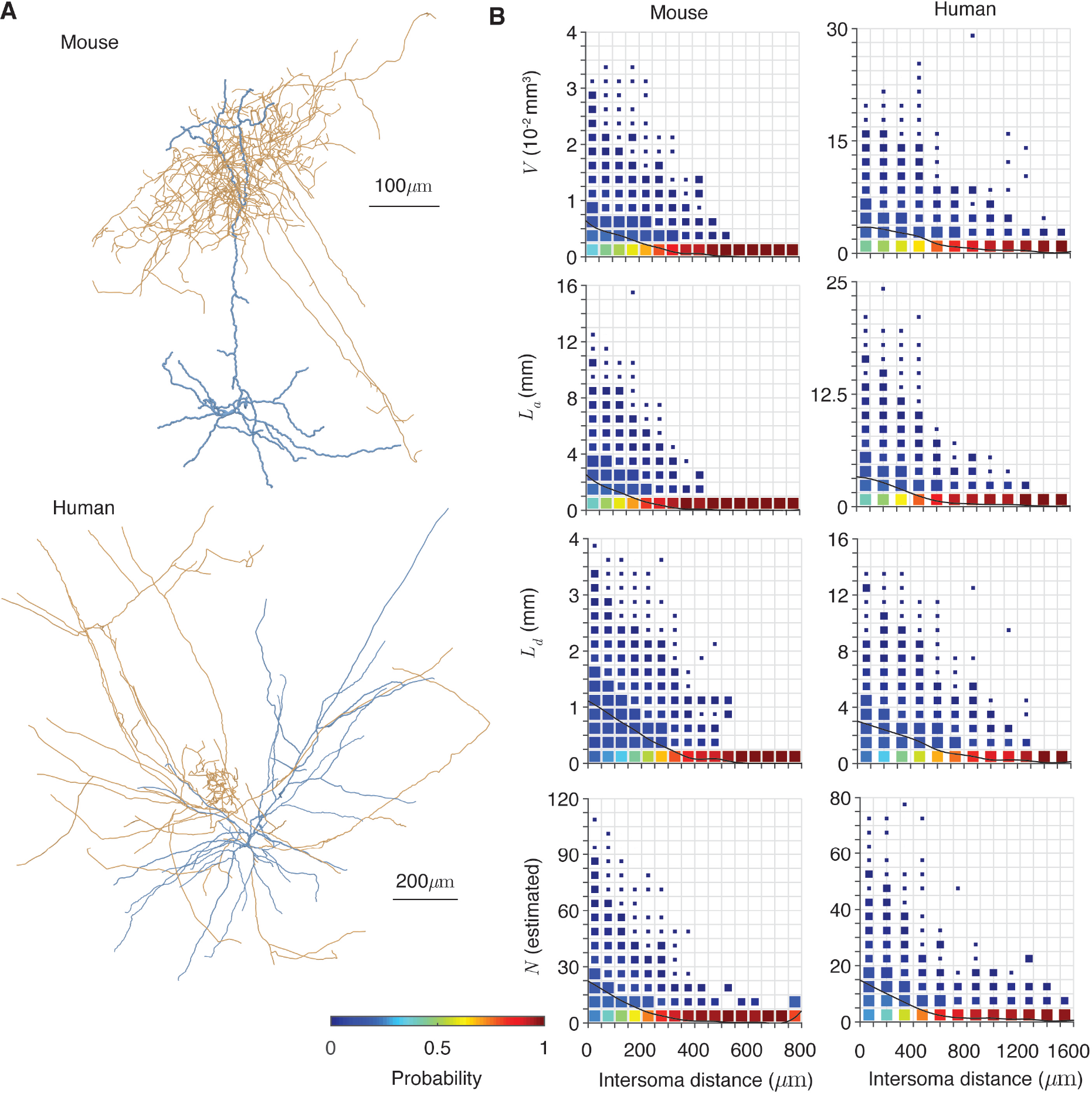
Applying the prediction of putative connectivity. **A** Example morphologies of axonal (orange) and dendritic (blue) trees from mouse (top) and human (bottom) cortex (Allen Brain Institute, 2015). Diameters are increased by 1*μ*m to increase visibility. **B** The dependencies of shared volume *V*, axonal length *L_a_*, dendrite length *L_d_*, and expected putative synapse number *N* on intersoma distance for mouse (left) and human (right) cells. 10^4^ random pairs are taken in each case. Distributions are normalised for each interval of intersoma distance and square sizes scale linearly with the occurrance of each probability in the grid. Black lines show the mean of each quantity.

## Discussion

### Summary

We have demonstrated that the expected number of putative synaptic contacts between a pair of neurons is given by a simple equation (Eq 1) and depends on the volume of axo-dendritic overlap, the lengths of axon and dendrite within this region, and the maximum length of dendritic spines. Moreover, if these four factors are controlled for, the number of synapses is insensitive to domain shape, intersoma distance, centripetal bias, and morphological pathologies that cause clustering. These results also hold for reconstructed cortical axons and dendrites at random distances and orientations and are simply applicable to real cells. When applying our results to cells reconstructed in context we found an excess of potential connectivity, meaning that each pair of cells could potentially be connected multiple times. The potential contacts are distributed in space and, on average for each cell class, each axon could be expected to target every available dendritic compartment, whether defined by branches or by electrotonic attenuation. This suggests that such cortical microcircuits require potential connectivity to specific dendritic compartments, rather than simply at a cellular level, and are able to achieve this.

### Context

Theoretical estimates of potential synaptic connectivity between pairs of neurons have long been sought alongside experimental approaches to clarify and generalise the principles observed. It is useful to consider the other approaches taken to this problem, to demonstrate how the estimates of our Eq 1 fit into the broader literature on putative connectivity.

An early analytical approach by Uttley (1955) estimated the number of close appositions of axon and dendrite by assuming random and independent neurite growth and its implications on the resultant densities of cable. Liley & Wright (1994) built on this, using a spherically symmetric simplification of dendritic density based on the Sholl intersection profile (Sholl, 1953) to estimate the probabilities of given numbers of anatomical connections forming between pairs of cortical cells. Stepanyants et al (2002) considered the relationship between potential and actual connectivity, introducing the concept of the filling fraction and modelling the number of potential contacts using the independent features of the axon and dendrite. Kalisman et al (2005) used the detailed structure of complete reconstructions to generate more realistic cylindrically symmetric estimates of cortical neurite density and find the probability of putative synapses forming between cell pairs. Stepanyants et al (2004) studied specific connectivity between interneurons and pyramidal cells, confirming that specificity in connections could arise from overlapping volumes of axon and dendrite even in the absence of correlations between neurite branches and finding a linear increase in potential connectivity with maximum spine distance. Chklovskii (2004) considered the volume of connected neuronal circuits and determined the relative contributions of branched axons, dendrites, and dendritic spines to allowing physiological levels of compactness. In particular, this study estimates the expected number of afferent synapses onto a given neuron from the length of its dendrite, maximal spine length, and the amount of axon in the surrounding tissue. Amirikian (2005) also assumed a cylindrically symmetric density of synapses (based on the dendritic density given by the Sholl intersection profile in a similar manner to Liley & Wright (1994)) and found good agreement with data for a number of cell classes, in particular correctly determining both axo-dendritic and axo-somatic synapses. Wen et al (2009) considered the connectivity repertoire, the number of possible patterns of synaptic contacts a given dendrite can receive. During this study, estimates of the numbers of synaptic contacts onto Purkinje and pyramidal cell basal dendrites were derived in terms of their length, maximal spine distance, and spanning field. Hill et al (2012) took a different approach, using full reconstructions to determine putative synaptic contacts and finding excellent agreement with the potential synapses found experimentally. van Pelt & van Ooyen (2013) directly compared the connectivity predictions from neurite density fields with those of the full arbors, finding that whilst predictions from density fields do generally match the expected number of anatomical contacts, they are unreliable estimators of both the absolute connection probability (the probability that at least one anatomical contact occurs) and the true distribution of possible contact numbers. McAssey et al (2014) greatly expanded on the previous study, simulating the connections between 10,000 artificially constructed dendrites and relating these to those derived from the neurite density fields. Reimann et al (2015) developed the approach of Hill et al (2012) using observed axo-dendritic overlaps to refine the connectivity patterns, producing a simulated cortical column connectome that was explored by Markram et al (2015). Aćimović et al (2015) used the gaussian description of dendritic density introduced by Teeter & Stevens (2011) to examine how changing the parameters of simplified dendrites could affect the connectivity structure of a large network.

Our approach can be seen as a broad generalisation of the above studies. The estimate of Eq 1 is not dependent on the local statistics of different neuronal types, such as those of Kalisman et al (2003, 2005), Hill et al (2012), and Reimann et al (2015, 2017), nor does it rely on the density of neurite within a spanning field such as the work of Uttley (1955), Liley & Wright (1994), Amirikian (2005), van Pelt & van Ooyen (2013), McAssey et al (2014), and Aćimović et al (2015). The simple form of Eq 1 is similar to the equations derived by Stepanyants et al (2002) and Chklovskii (2004) for the total number of synapses in a volume of neural tissue; the novelty of our result is that such a simple equation can be reliably applied to individual cell pairs.

### Outlook

This study provides a simple way to verify the degree of targeting within a neuronal circuit; for a given pair of neurons, the expected number of putative synaptic contacts is predictable by our equation. When the number of close appositions is consistently different from that predicted here, there is evidence of local targeting in the neurite growth processes and potential for a higher degree of hard-coding or specificity in a circuit. This could help to explain some of the controversy over the applicability of versions of Peters’ rule in different systems (Rees et al, 2017), and will become more valuable as connectome data becomes increasingly available

We would also like to highlight the importance of the location and shape of the axo-dendritic overlap. Our results suggest that this is both important for the overall connectivity estimate and sufficient to predict the putative locations of inhibitory connections that are known to be highly specific (Ascoli et al, 2008). Hill et al (2012) found that the statistical description of neuronal morphologies in isolation failed to predict the connectivity of such interneurons, in particular chandelier cells where the interaction of axon and dendrite is assumed to provide additional growth guidance. By taking the overlapping volume relative to the two cells, this factor is already accounted for and a random distribution of neurite within this volume is sufficient to predict connectivity. This contributes to the recognition of the importance of neurite spanning fields in the context of neighbouring cells as a powerful first step towards understanding the structure and function of a given cortical system (Sümbül et al, 2014; Bird & Cuntz, submitted).

We should note again that putative synaptic contacts are not necessarily bridged by actual dendritic spines, and data suggests that the ratio of actual to putative contacts, the filling fraction, is often very low (Jiang et al, 2015; Kasthuri et al, 2015; Lee et al, 2016). It should not even be assumed that the filling fraction is consistent between different cell pairs as synaptic existence and effective weights will depend on network inputs (Hebb, 1949; Yu et al, 2009; Lee et al, 2016; Ferrarese et al, 2018), intrinsic electrophysiology (Llinas, 1988; Ascoli et al, 2008), and variable short- (Zucker & Regehr, 2002) and long-term (Le Bé & Markram, 2006; Betley et al, 2009; Markram et al, 2012) synaptic plasticity. There is also no direct correspondence between the number of functional anatomical contacts and synaptic weight due to variability in postsynaptic spine size and sensitivity (Arellano et al, 2007; Bhumbra & Beato, 2013) as well as presynaptic vesicle number and release probability (Loebel et al, 2009; Bird et al, 2016). Nevertheless, the close appositions between axon and dendrite analysed here do provide a stable structural backbone (Trachtenberg et al, 2002; Knott et al, 2002; Chow et al, 2009), being in the first place necessary for any connectivity at all and allowing neuronal activity to dynamically reshape functional connectivity on timescales from the tens of milliseconds of short-term depression to the years and decades of long-term plasticity (Hebb, 1949; Zucker & Regehr, 2002). The recent review of different interpretations of Peters’ Rule by Rees et al (2017), notes that a number of experimental papers reject the hypothesis that functional contacts form at random in cortex (Potjans & Diesmann, 2014; Kasthuri et al, 2015; Lee et al, 2016). Each of these papers does however find that close axo-dendritic appositions do appear to occur randomly and this is not a contradiction of our hypothesis here. In other neuronal systems, such as the retina (Kim et al, 2014) or fly brain (Takemura et al, 2015), neurite interactions do appear to be more hardwired than in cortex or hippocampus, so caution would be necessary when applying these findings elsewhere.

The excess putative connectivity we observe in the cortical dataset is in line with the results of Kasthuri et al (2015) and Lee et al (2016), but higher than the earlier estimates cited in Stepanyants et al (2002). The earlier paper interpreted a low filling fraction (ratio of actual to potential contacts) as a signature of a more complex system as more patterns of connectivity can be implemented by choosing a selection of available contacts, enhancing the information storage capacity of the synapse. This view does not fundamentally conflict with that presented here, rather we highlight that the absolute number of potential contacts is potentially very high and show how this could allow for the implementation of connectivity to specific dendritic compartments. The location of dendritic inputs relative to one another are already known to be key to the response properties of neurons (Mel, 1993; Poirazi et al, 2003; Polsky et al, 2004; Behabadi et al, 2012; Gidon & Segev, 2012); and it is unsurprising that neurite structure allows for specific input patterns to be implemented.

Neurites allow a broad range of connectivity patterns (Wen et al, 2009; Markram et al, 2015; Jiang et al, 2015), whilst obeying optimality principles in volume, length, and signal delays (Chklovskii, 2004; Budd et al, 2010; Cuntz et al, 2010). The specific layout of axonal inputs and branching principles can combine to create diverse dendritic shapes (Cuntz, 2012; Cuntz et al, 2012), which in turn lead to the connectivity patterns seen in neuronal circuits (Hill et al, 2012; McAssey et al, 2014; Potjans & Diesmann, 2014). Whilst the design principles leading to optimal connectivity and optimal wiring could appear at odds, both approaches have proved successful in reconstructing the major features of real neurons (Braitenberg & Schüz, 1998; Stepanyants & Chklovskii, 2005; Cuntz et al, 2007; Wen et al, 2009; Cuntz et al, 2010); this study reaffirms that optimal design principles are common between both goals. Overall our work provides an intuitive way to estimate the putative synaptic connectivity of microcircuits, greatly simplifying the parameters necessary for analytical and numerical studies of the structure of biophysically detailed neuronal networks, including at the level of dendritic compartments.

## Acknowledgments

We acknowledge funding through BMBF grant 01GQ1406 (Bernstein Award 2013) to HC.

## Materials and methods

### Data and algorithm availability

All simulations were done in MATLAB (ver. 2018b) using the TREES toolbox (ver. 1.15) and custom scripts. The morphologies used in the paper were downloaded from NeuroMorpho.Org (www.neuromorpho.org, Ascoli et al (2007)) and the Cell Types database of the Allen Institute for Brain Science (Allen Brain Institute, 2015) with sources indicated in Table 3. Selection of cells and preprocessing for the morphologies in Figure 5 were performed using the Allen Software Development Kit (freely available at http://alleninstitute.github.io/AllenSDK/install.html) in a Jupyter Notebook (ver. 5.5) running Python (ver. 3.7). All data and code necessary to reproduce the figures and tables in the paper are included as Supplementary File 1.

**Table 3.** Table of data sources and morphologies shown in figures. For the first two rows NeuroMorpho accession numbers (NMOs) are stated, for the last two rows Allen Cell Types Database IDs are stated.

| Dataset | Figures & Tables | Source | Example morphologies |
| --- | --- | --- | --- |
| Rat barrel cortex<br>75 cells | Figure 3<br>(panels a to e) | Marx et al | Fig 3a - Axon NMO.64655,<br>Dendrite NMO.64634 |
| Mouse visual cortex<br>323 cells | Figure 4*, Table 2<br>*(all panels except f) | Jiang et al | 4a - BTC NMO.61572,<br>BTC NMO.61573, L5MC NMO.61669,<br>ChC NMO.61626, L5MC NMO.61670,<br>BPC NMO.61596, ChC NMO.61627 |
| Mouse cortex<br>126 cells | Figure 5<br>(all parts) | Allen | 5a - Axon Cell ID:584236936,<br>Dendrite Cell ID:584872371 |
| Human cortex<br>112 cells | Figure 5<br>(all parts) | Allen | 5a - Axon Cell ID:566350399,<br>Dendrite Cell ID:527941296 |

Of particular general utility are the following new MATLAB Trees Toolbox functions:

- `dscam_tree` - Applies the DSCAM null algorithm (described below) to a tree to produce clustered dendrites.
- `M_atten_tree` - Estimates the number of dendritic compartments based on electrotonic attenuation.
- `peters_tree` - Determines the number of putative anatomical contacts between two neurites.
- `share_boundary_tree` - Determines the boundary of the overlap of two neurites and their respective lengths within this region.

### Generalised minimum spanning trees

Synthetic neurites are produced using the generalised minimum spanning tree (MST) algorithm described by Cuntz et al (2010) and available as part of the Trees Toolbox package for MATLAB. Typically neurites are simulated by uniformly randomly distributing a number of points in a cube with sides of length 200*μ*m and connecting them into a generalised MST using the Trees Toolbox function `MST_tree`. The number of points for both the axonal and dendritic trees is typically taken to be 65, the trees are generated in cubes with 25*μ*m between their centres and the balancing factors of the axonal and dendritic trees are 0.7 and 0.2 respectively.

In order to systematically vary the parameters *L*_*a*_, *L*_*d*_, and *V* it is necessary to vary the numbers of points used to generate the axons and dendrites, the separation of the domains, and the scale of the neurons. To allow for a consistent search of parameter space, we use the fact that any intermediate step in the construction of a minimum spanning tree by a greedy algorithm forms the minimum spanning tree of the set of points it connects (Prim, 1957). This allows the Matlab `patternsearch` optimisation function to be used to find pairs of trees that have the desired properties. Optimisation is stopped when the deviation from the desired properties is less than 5% in total. Such optimisations are potentially time consuming and the Neuroscience Gateway cluster was used to test different approaches (Sivagnanam et al, 2013). Figure S1 plots the resultant distributions of *L*_*a*_, *L*_*d*_, and *V* used to construct Figure 1.

### Definining the axo-dendritic overlap

To define the boundary of the axo-dendritic overlap (Fig 1a), the following procedure was used. The boundaries of the axonal and dendritic arbors were computed using the Trees Toolbox `boundary_tree` function (Cuntz et al, 2010; Bird & Cuntz, submitted). Both arbors were resampled to a resolution of 1*μ*m using the `resample_tree` function. The set of axonal nodes lying within the dendritic boundary and the set of dendritic nodes lying within the axonal boundary were selected. A boundary was constructed around this set of points using the mean convexities of the two neurite arbors (see below).

### Boundaries and convexities

Boundaries are constructed using *α*-shapes (Edelsbrunner et al, 2006). An *α*-shape is a generalisation of the convex hull of a point set whereby a boundary is a set of simplices (triangles in three dimensional space) constructed by placing balls of radius 1/*α* over the point set so that all points are contained within the ball and the vertices of the bounding simplex lie on the surface of the ball. To enable this construction, *α* must lie between 0 and some small positive value (the generalisation to negative values is not necessary here), but small changes in *α* do not necessarily lead to distinct *α*-shapes. An *α*-spectrum is constructed as the set of *α*-intervals which define distinct boundaries and a parameter known as the shrink factor defines the proportion of the way through this spectrum that an *α* value is chosen. In practice this procedure is implemented through the MATLAB boundary function. The shrink factor is taken as one minus the convexity of a tree (Bird & Cuntz, submitted).

To define the convexity of a tree, we take the set of terminal neurite points and see what proportion of the direct paths between them lie entirely within the tightest boundary (a shrink factor of 1) that contains all termination points. For a convex hull, all such paths would lie within the boundary and so this measure gives the relative difference between the two extreme shrink factors. We refer to this value, between 0 and 1 as the convexity (where 1 gives an entirely convex neuron). The Trees Toolbox function `convexity_tree` carries out this computation (Cuntz et al, 2010; Bird & Cuntz, submitted).

### Estimation of putative synapses

Putative synaptic contacts are identified by the close apposition of dendrite and axon. An algorithm to do this for neurites in the Trees Toolbox format has been developed. The first step is to resample neurites into sections of a consistent length; this is taken to be 1*μ*m, although shorter distances may be appropriate if the neurites have greater tortuosity. This is done with the existing `resample_tree` function. The second step is to identify the pairs of resampled axonal and dendritic nodes that are less than the maximal spine distance *s* apart. This will typically lead to a large number of pairs of axonal and dendritic nodes that are very close together due to the connected structure of each neurite. It is rare to find such an arrangement in the data, as distinct synaptic contacts between a pair of neurons are typically more than a few microns apart. To recover a realistic distribution of contact locations and numbers, a greedy deletion step is applied. The closest pair of axonal and dendritic nodes are chosen as a synaptic contact. Then all putative contacts where the axonal node is within a 3*μ*m distance of the axonal node in the existing contact and the dendritic node is within a 3*μ*m distance of the dendritic node in the existing contact are deleted. The closest remaining pair of axonal and dendritic nodes is chosen as a second synaptic contact and the deletion step is repeated until all inappropriate contacts are removed.

### Derivation of analytical estimate

The expected number of putative contacts can be derived by considering the number of crossings of a cylinder of radius *s* centred on the dendrite by axonal branches. We follow Stepanyants et al (2002) and consider axons and dendrites as a set of isotropically distributed straight segments that are significantly longer than the maximum spine distance *s*. Let an axonal segment of length 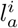 make an angle *θ*_*i,j*_ with a dendritic segment of length 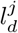 both lying within the volume *V* . Then the probability *p*_*θ*_ that they intersect is

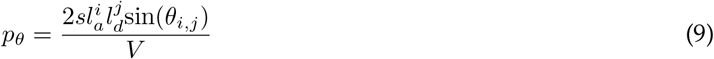

Taking a sum over *i* and *j* gives the expected number of synapses within the volume in terms of the angles *θ*_*i,j*_. Assuming that the lengths of neuritic segments are independent allows the sum to be separated as

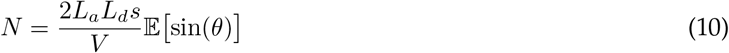

As the segments are assumed to be distributed isotropically, the expected value of sin(*θ*) is 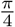 and this leads directly to Eq 1.

### Dimensions of dendritic domains

The three additional domains in Figure 2a are chosen to match the volume of the cube with sides of 200*μ*m. Therefore the sphere has radius 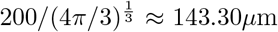 and the soma is located in the centre. The cylinder is chosen to have the same height and cross-sectional diameter, both are 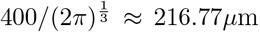, again the soma is positioned in the centre. The cone is chosen to have equal height and terminal diameter, both are 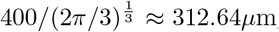; in this case the soma is located at the point of the cone. It should be noted that the relationship between the volume spanned by a dendrite and the volume in which target points are distributed are not the same; the former will be bounded above by the latter as the number of target points approaches infinity (Ripley & Rasson, 1977). The volume bounded by a generalised minimum spanning tree within a given target domain depends on the shape of that domain, the balancing factor, and the soma location. Figure S2a shows the relationship between dendrite length and volume in the different domains.

### DSCAM null model

DSCAM null mutants have unusually clustered dendrites (Schmucker et al, 2000; Soba et al, 2007). Such dendrites do not fill space efficiently and are not well-reproduced by the standard generalised MST model of Cuntz et al (2010). To implement an algorithm to produce realistic synthetic dendrites, the following iterative procedure was defined:

1. A node *a* on the tree is uniformly randomly chosen.
2. The closest node *b* that is not directly connected to the first node *a* and is more than 2*μ*m away from it is identified.
3. The first node *a* is moved 10% closer to node *b*.
4. Steps 1-3 are iterated K times.

This algorithm produces dendrites with clustered branches that resemble those of DSCAM null mutants. The restriction of a minimal 2*μ*m distance in step 2 is necessary to prevent branches becoming ‘paired’ and merging together so that additional steps of 10% of the distance between them do not alter the geometry of the tree. It should also be noted that this algorithm is best applied directly to the output of the existing Trees Toolbox `MST_tree` function without any resampling. Increasing the number of steps K qualitatively changes the relationship between dendrite length and the volume spanned (Fig 2d). For the DSCAM null synthetic dendrites in Figure 2, K = 100, 400, and 700.

### Reconstructed morphologies

We retrieved 75 neurons from NeuroMorpho.org (Ascoli et al, 2007) to apply our algorithm to reconstruction data. The rat barrel cortex neurons were originally obtained by Marx et al (2017). The data set included five different neuron types (pyramidal-like, multipolar, horizontal, tangential and inverted) from layer 6b (N = 49) and SP (N = 26). The reconstructions were preprocessed in the following way: first they were resampled to 1*μ*m line pieces. To separate dendrites and axons, all nodes that were not labelled as axon were deleted and the remaining nodes saved as a dendrite. The axons were obtained in the same way, but with soma and dendrite removed. The volumes and cable lengths differed widely between the reconstructions (Fig S3). Then dendrites and axons were randomly paired and the axon was shifted random amounts between 0 to 100*μ*m in the X, Y, Z directions and rotated uniformly randomly. A resulting example pair of axon and dendrite can be seen in Fig 3a. The number of estimated and measured putative synaptic contacts were calculated for 10,000 of these combinations.

### Distribution of *n*

Assuming that putative contacts are formed uniformly and independently means that the distribution of the number of contacts is Poisson

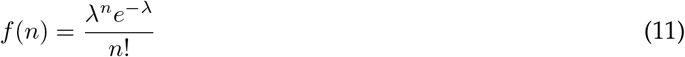

The parameter *λ* gives the mean and variance of the distribution, which must be the same. A common generalisation of the Poisson distribution to allow for these moments to differ is the Pólya distribution *g*(*n*) (DeGroot & Schervish, 1983).

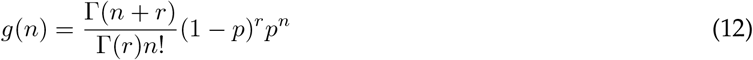

where 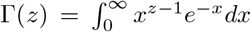 is the gamma function. The mean and variance are given by *pr/*(1 − *p*) and *pr*/(1 − *p*)^2^ respectively. To fit the connection probability *p*_*c,g*_ to 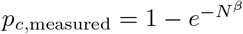 (where *β* = 0.5437), the parameters *p* and *r* obey

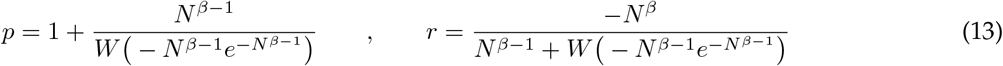

where *W* is the Lambert *W* function satisfying *z* = *W*(*z*)*e*^*W*(*z*)^ for any complex number *z*. These parameters give a poor match to the measure variance for small values of *N* (blue dashed line in Figure 3d).

The skewness *γ* of a random variable *X* is defined as 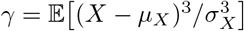 and measures the degree to which probability mass lies above the mean. Positive values mean that more mass lies above the mean and negative that more mass lies below; further intuitions about the meaning of this statistic can often be flawed (von Hippel, 2005). The unbiased skewness *γ*_*x*_ of a set of samples {*x*_*i*_}_*i*=1,2,..,*k*_ of size *k* with sample mean *x̄* is given by

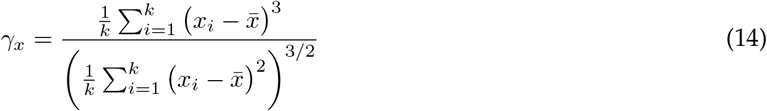

The skewness of the values of *n* for each interval of *N* in Figure 3 are plotted in Supplementary Figure S3c. The skewness is well-described by a function of the form *γ*(*N*) = *e*^−*cN*+*d*^ where the fitted parameters (with 95% confidence intervals are) *c* = 0.1184 (0.0860, 0.1509) and *d* = 0.7615 (0.6268, 0.8962). This is shown by the black line and shaded region in Figure S3c. The skewnesses of the Poisson (*γ*_*f*_) and Pólya (*γ*_*g*_) models are given by

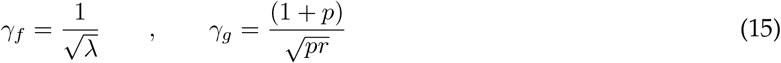

Plotting these values with parameters fitted to the measured mean and variance (in the case of the Pólya distribution) shows a poor match to the measured skewness (blue and red lines in Figure S3c), particularly for smaller values of *N*.

The Pólya distribution can be considered a generalisation of the negative binomial distribution, which counts the number of Bernoulli successes at constant probability *p* before *r* failures occur, to allow for a non-integer value of the stopping parameter *r*. Similarly, the negative hypergeometric distribution gives the number of Bernoulli successes that occur before a certain number of failures when each success changes the probability of future successes. The usual interpretation is drawing from a finite set without replacement. Making a similar generalisation to non-integer parameters, the negative hypergeometric distribution *h*(*n*) is given by

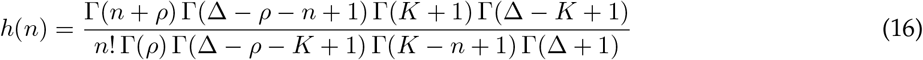

The mean *μ*_*h*_ and variance 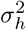 are given by

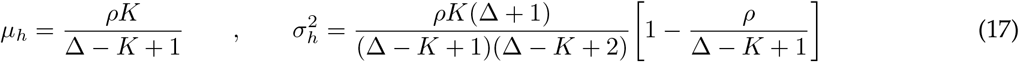

The interpretation of the parameters is typically that *n* counts the number of Bernoulli successes before *ρ* failures occur; initially the probability of success is *K/*Δ, but each success decreases (and each failure increases) the subsequent success probability. The parameters therefore obey *K* ≤ Δ and *ρ* ≤ Δ − *K*. There is no closed-form expression for the skewness so we obtain it numerically (see below).

Fitting the three central moments, *μ*_*h*_, 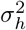, and *γ*_*h*_, as well as *p*_*c*,measured_ to the data for each value of *N* gives the curves in Figure S3d. The mean is fitted exactly and the sum of the relative differences of the other three statistics from their true values is minimised using the Matlab `fmincon` function.The negative hypergeometric distribution is a good fit for the observed distributions of *n* for small values of *N*. Figures S3g and h show the best fits of the Poisson, Pólya, and negative hypergeometric distributions to the observed distributions of *n* for *N* = 2, and 5.

As *N* grows, the negative hypergeometric and Pólya models fitted in this way diverge. Figure S3e shows the Kullback-Leibler divergence of Eq 16 from Eq 12 (Kullback & Leibler, 1951). The Kullback-Leibler divergence *D*_*KL*_(*H*‖*G*) quantifies the information gain (in nats) from using a distribution *h*(*n*) instead of *g*(*n*) and is defined as

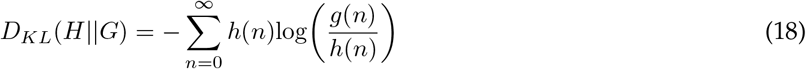

This approaches zero for *N* ≈ 10 before growing; using the more complex Eq 16 instead of Eq 12 has little additional utility above this value. The confidence intervals in Table 2 are therefore calculated using the most appropriate distribution.

Figure S3i plots the distributions of *n* for each integer value of *N* under the negative hypergeometric model.

### Numerical evaluation of Eq 16 and estimation of skewness *γ*_*h*_

To evaluate Eqs 12 and 16 for large values of the parameters or *n* requires a numerical approximation to the gamma function 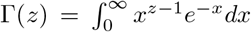 for large arguments *z*. Following an approach in Bird et al (2016), Stirling’s approximation allows the gamma function to be evaluated as

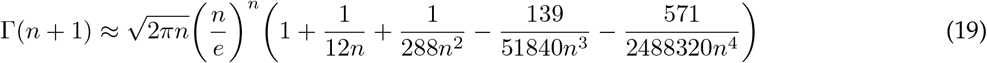

In particular, this form allows the logarithm of each factor to be computed separately and so gives an accurate result for the probability mass when large terms in the numerator and denominator of Eq 16 cancel out. The skewness *γ*_*h*_ is computed using the forms of *μ*_*h*_ and 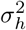 in Eq 17 and the non-central moment 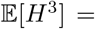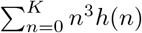 evaluated over the range where *n*^3^*h*(*n*) is above the smallest IEEE positive double precision number (≈ 2.23 × 10^−308^). Hence

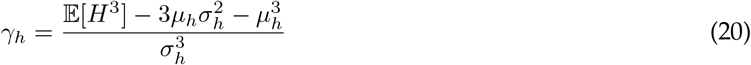

### Reconstructed microcircuits

We retrieved 323 neurons from NeuroMorpho.org (Ascoli et al, 2007) to apply our algorithm to reconstruction data in context. The mouse visual cortex neurons were originally obtained by Jiang et al (2015). Our dataset contained many fewer morphologies than those reported in the original paper, but comprised the full set of publicly available complete reconstructions from a slice with more than one cell at the time of writing. The data set included sixteen different neuron types (see Table 2) from cortical layers I, II/II and V. Cell-type labels follow directly from the labelling in the initial paper using the approach of Sümbül et al (2014) where classification is done on the distribution of neurite within the cortical layers. The abbreviations used in Figure 4 and Table 2 are as follows: SBC-like single-bouquet cell like, eNGC extended neurogliaform cell, L23MC layer 2/3 Martinotti cell, L23NGC layer 2/3 neurogliaform, BTC bitufted cell, BPC bipolar cell, DBC double-bouquet cell, L23BC layer 2/3 basket cell, ChC chandelier cell, L23Pyr layer 2/3 pyramidal cell, L5MC layer 5 Martinotti cell, L5NGC layer 5 neurogliaform cell, L5BC layer 5 basket cell, HEC horizontally extended cell, DC deep-projecting cell, L5Pyr layer 5 pyramidal cell.

The total number of neuron pairs (where pairs are imaged from the same slice) was 434, of which 233 had more than one putative contact. The reconstructions were preprocessed in the following way: first they were resampled to 1*μ*m pieces. To separate dendrites and axons, all nodes that were not labelled as axon were deleted and the remaining nodes saved as a dendrite. The axons were obtained in the same way, but with soma and dendrite removed. The volumes and cable lengths differed widely between the reconstructions (Fig sS4a and b).

### Nearest- and all-neighbour ratios

To establish the spatial distribution of potential contacts within the shared volume, we considered both the nearest- and all-neighbour ratios (Chandrashekhar, 1943). The nearest-neighbour ratio quantifies the pairwise spatial correlation of points in space, whereas the all-neighbour ratio seeks to capture higher order correlation structure. The nearest-neighbour ratio of a set of points bounded by a given volume is defined as the mean ratio of nearest-neighbour distances to that of all possible sets of uniformly distributed points with that same space. The all-neighbour ratio in contrast is the mean distance of each point to the centroid of all other points in the same space. A nearest-neighbour ratio of close to one implies that points are distributed uniformly in space, whereas an all-neighbour ratio of one implies that at larger scales points are uniformly distributed. For each overlapping volume with *n* potential contacts, we generated 10^6^ sets of *n* uniformly randomly distributed points to calculate the mean nearest- and all-neighbour distances. This also allowed the *p*-values to be computed from the proportion of random point sets that have a more extreme ratio in a given volume than the measured synaptic sites. For our data, the nearest- and all-neighbour ratios are highly correlated (Fig S4c), suggesting that the higher-order correlation structure is caught by the pairwise, local measure.

The neighbour ratios can often be applied in a biased manner, particularly when the boundary within which the points are to be distributed is defined by the set of points themselves (Ripley & Rasson, 1977; Anton-Sanchez et al, 2018). Our application, with the boundary defined by the overlapping neurite arbors, rather than strictly the synaptic locations should be relatively free of this bias. To check whether this is indeed the case we examined whether the volumes spanned by the boundary around the putative contacts differed systematically from the boundary around all uniform random point sets with the same number of members. Figure S4d shows that this is not the case.

### Dendritic compartments

The passive electrotonic properties of each interneuron type were estimated from the somatic input resistance values reported in Jiang et al (2015) (Tables S1 S2) and those of the pyramidal cells from the values reported by Guan et al (2015) (Figure 2). There will always be a continuum of pairs of values of axial resistivity *r_a_* and membrane conductivity *g*_*m*_ that lead to a given somatic input resistance for a given morphology; different pairs will also typically lead to slightly different electrotonic response properties elsewhere in neurons with tapering dendrites due to the distinct contributions of *r*_*a*_ and *g*_*m*_ to the local electrotonic length constant (Goldstein & Rall, 1974; Bird & Cuntz, 2016). We therefore restricted the allowed values to a physiological range, with axial resistivity between 50 and 200 Ωcm and membrane conductivity allowed to vary between 0 and 10^−3^ Scm^−2^. The MATLAB patternsearch optimisation function was used to minimise the difference between the mean somatic input resistance for each class of morphologies and that reported experimentally. The starting values were of *r_a_* = 100 Ωcm and *g*_*m*_ = 5 × 10−5 Scm^−2^ and the optimisation was stopped once the difference between means was below the standard error quoted in the above papers. In practice, the axial resistivities remained at their initial value in all cases. The somatic input resistances reported by Jiang et al (2015) and Guan et al (2015) and the resultant membrane conductivities estimated by our algorithm are tabulated in Supplemental Table 1. The membrane conductivities derived in this way are higher than is sometimes reported for mouse cortical neurons in detailed electrophysiological studies, but fit the reported somatic input resistance and are consistent with the relatively fast membrane time constants also reported (Gentet et al, 2000).

The electrotonic signature, the set of transfer resistances from each node to all other nodes, was computed using the existing Trees Toolbox function `sse_tree`. The compartmentalisation was defined by assigning a threshold value, in this case 0.13995, and finding the regions of the tree within which voltages do not attenuate below this proportion of the maximum input resistance. Different thresholds lead to different numbers of compartments in a given dendrite (Fig S4f).

### Compartments containing a synapse

The expected number of synaptic contacts necessary to innervate *M* dendritic compartments (*N*_Complete_ in Eq 6) is given by the solution to a standard problem in probability: the coupon collector’s problem (Blom et al, 1993). The standard statement of (the simplest case of) this problem is that given a set containing elements distributed equally amongst *k* different classes, how many draws (independent and with replacement) are expected to be necessary until at least one element from each set has been drawn. This is equivalent to the problem of distributing contacts between dendritic compartments.

The distribution of compartments innervated by at least one of the *N* synaptic contacts (Eq 7) does not appear to be a widely-reported result, so we give a brief derivation here. Write 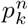 for the probability that *k* distinct compartments have been innervated by *n* contacts. Then for *n* = 1, a single contact must innervate a single compartment, hence 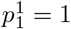 and 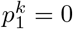 for all other values of *k*. For *n* > 1, 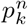 obeys the recursion relation

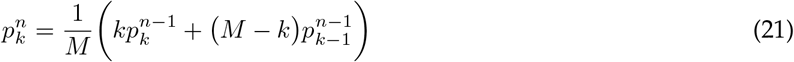

where the first term in brackets is the probability that the *n*-th contact innervates one of the *k* compartments already containing a contact and the second term is the probability that it innervates a new compartment. Solving the above difference equation leads to Eq 7.

### Reconstruction data to demonstrate the predictions

We retrieved 126 mouse and 112 human cortical neuron reconstructions from the Allen Brain Institute (Allen Brain Institute, 2015). Morphologies were chosen if they were marked as having a full dendrite and at least 200*μ*m of axon. Our dataset comprised the full set of publicly available reconstructions satisfying theses criteria at the time of writing. The data set included numerous different cell types. The reconstructions were preprocessed in the following way: first they were resampled to 1*μ*m pieces. To separate dendrites and axons, all nodes that were not labelled as axon were deleted and the remaining nodes saved as a dendrite. The axons were obtained in the same way, but with soma and dendrite removed. The volumes and cable lengths differed widely between the reconstructions (Fig S5, left hand panels). The cortical depths were recorded in the original dataset (Fig S5 for densities of neurite and somata with cortical depth). To generate the data for Figure 5, random pairs of axon and dendrite were chosen and displaced randomly uniformly by up to 125*μ*m in either direction in the plane parallel to both the slicing direction and cortical surface and by up to 25*μ*m in either direction in the plane perpendicular to the slicing direction and parallel to the cortical surface. Depth measurements are assumed reliable, but we introduce a small amount of jitter by displacing the depth by a normally distributed amount with mean 0*μ*m and standard deviation 10*μ*m. As these deviations are small compared to the range of possible cortical depths (Fig S5, right hand panels), much of the intersoma distance in Figure 5 comes from the layered structure of the cortex.

## Supporting information

**S1 File.** MATLAB code to reproduce all figures and data.

**Fig S1.**
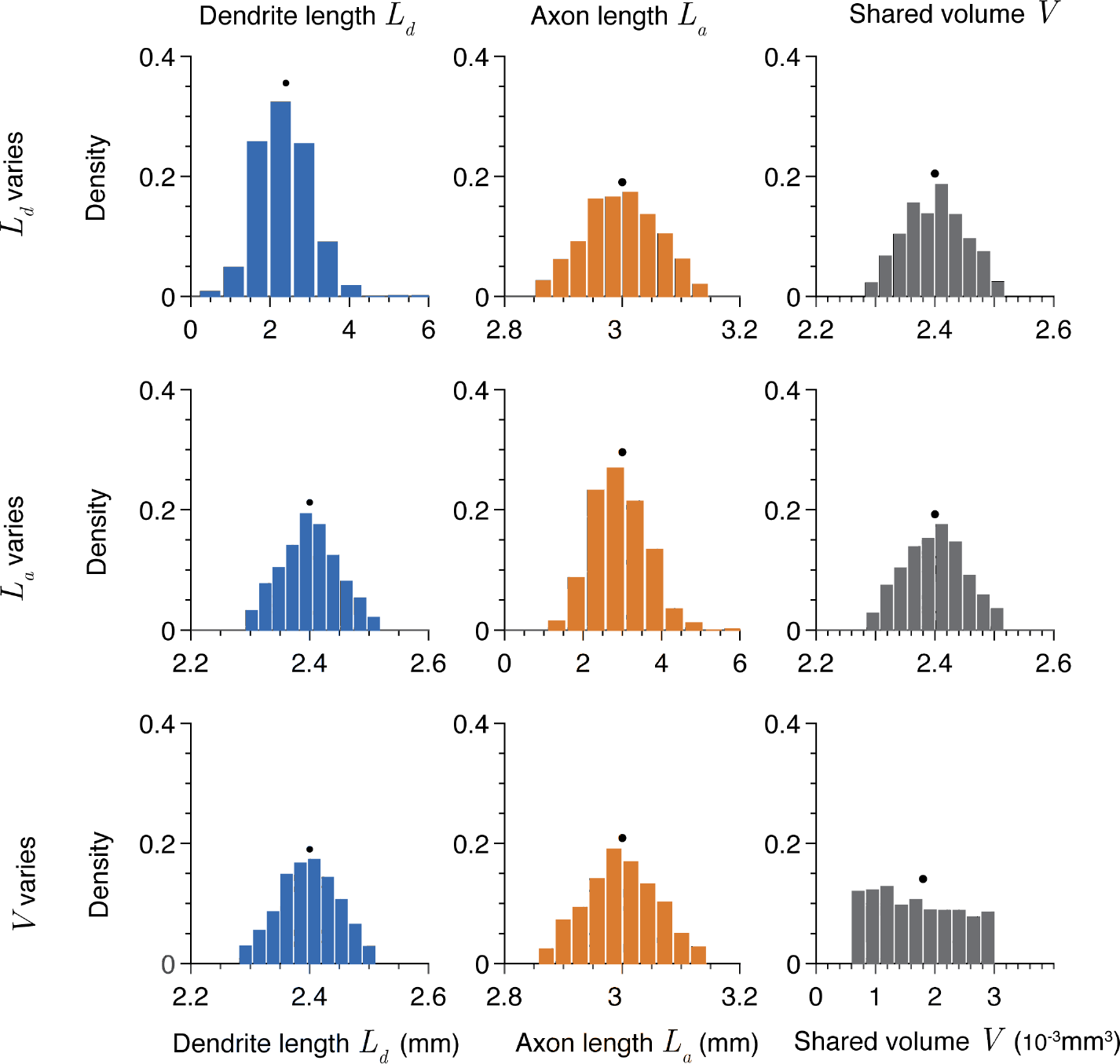
Supplement to Figure 1: Distributions of neurite length and shared volume for Figure 1. Distributions of dendrite length (left), axon length (centre), and shared volume (right) when dendrite length (top row), axon length (middle row), and shared volume (bottom row) vary. Generally, values are constrained to approximately *L_d_* = 2.4mm, *L_a_* = 3mm, and *V* = 2.4 × 10^−3^mm^3^ (shown by black dots).

**Fig S2.**
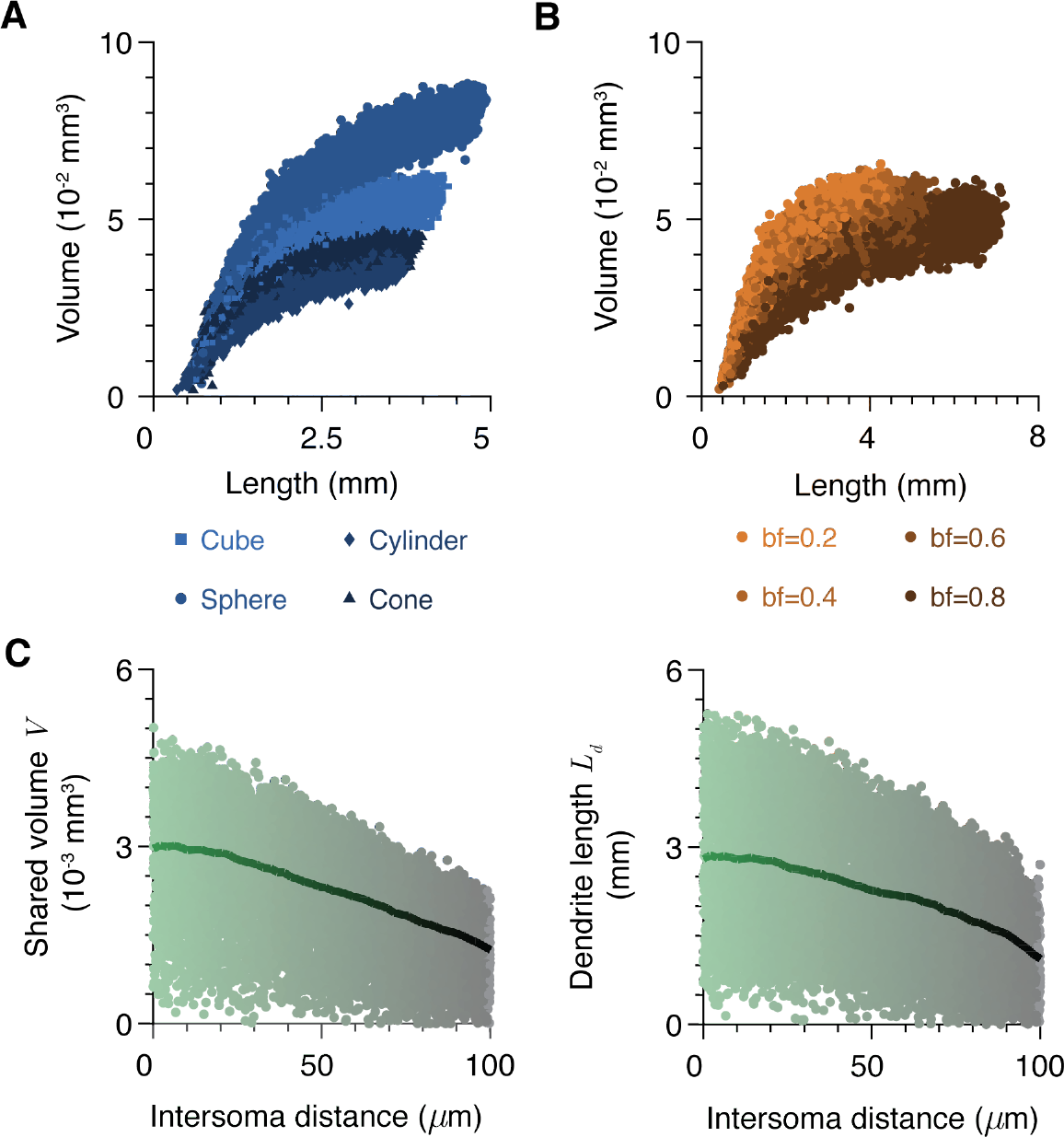
Supplement to Figure 2: Length and volume for dendrites for Figure 2 with different shapes and balancing factors and correlations with soma separation. **A** Dendrite spanning volume as a function of length for trees from Figure 2A bounded by a cube, sphere, cylinder, and cone. **B** Dendrite spanning volume as a function of length for trees from Figure 2B with balancing factors of 0.2, 0.4, 0.6, and 0.8. **C** Plots of shared volume (left) and dendrite length (right) as a function of intersoma distance for different morphologies (trees from Figure 2D).

**Fig S3.**
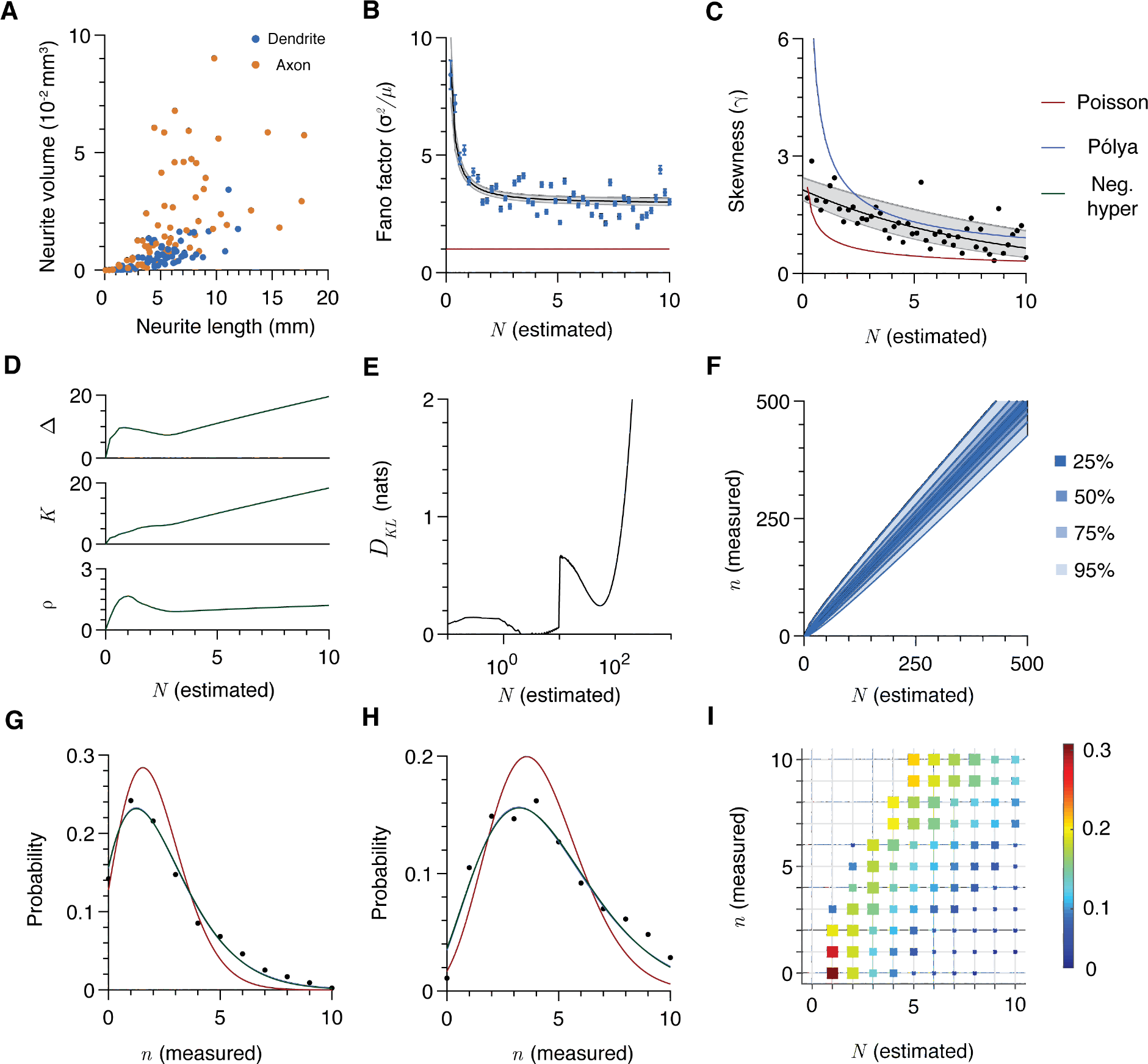
Supplement to Figure 3: Length and volume distributions for the reconstruction data in Figure 3 and parameters for the distribution of putative connection numbers. **A** Neurite spanning volume as a function of neurite length for the dendritic (blue) and axonal (orange) trees. **B** Fano factor (variance divided by mean) of the number of measured putative connections *n* as a function of the expected number *N* . The black line shows the fit of the variance equation (*aN + N ^b^*) and the grey shaded region shows the 95% confidence interval. The red line shows the Fano factor of the Poisson model. **C** Skewness of the number of measured putative connections *n* as a function of the expected number *N* . The black line shows the fit of the skewness equation (*e*^−*cN*+*d*^) and the grey shaded region shows the 95% confidence interval. Red and blue lines show the skewnesses of the Poisson and Pólya.models (Eq 15). **D** Parameters Δ, *K*, and *r* of the negative hypergeometric distribution (Eq 16) as a function of expected putative contact number *N* . **E** Kullback-Leibler divergence *DKL* (Eq 18) of the negative hypergeometric distribution (Eq 16) from the Pólya distribution (Eq 12) as a function of *N* . F Confidence intervals (25%, 50%, 75%, and 95%) for values of *n* as a function of *N* under the negative hypergeometric (for *N* < 10) and Pólya (for *N ≥* 10) models (Eq 16 and 12 respectively). **G** Example fit of Poisson (Eq 11, red line), Pólya (Eq 12, blue line), and negative hypergeometric (Eq 16, green line) models to data for *N* = 2. **H** As in **G** for *N* = 5. **I** Probability distribution of putative contact numbers *n* for each integer estimated putative contact number *N* under the negative hypergeometric model (Eq 16). Square sizes scale linearly with the occurrence of each probability in the grid.

**Fig S4.**
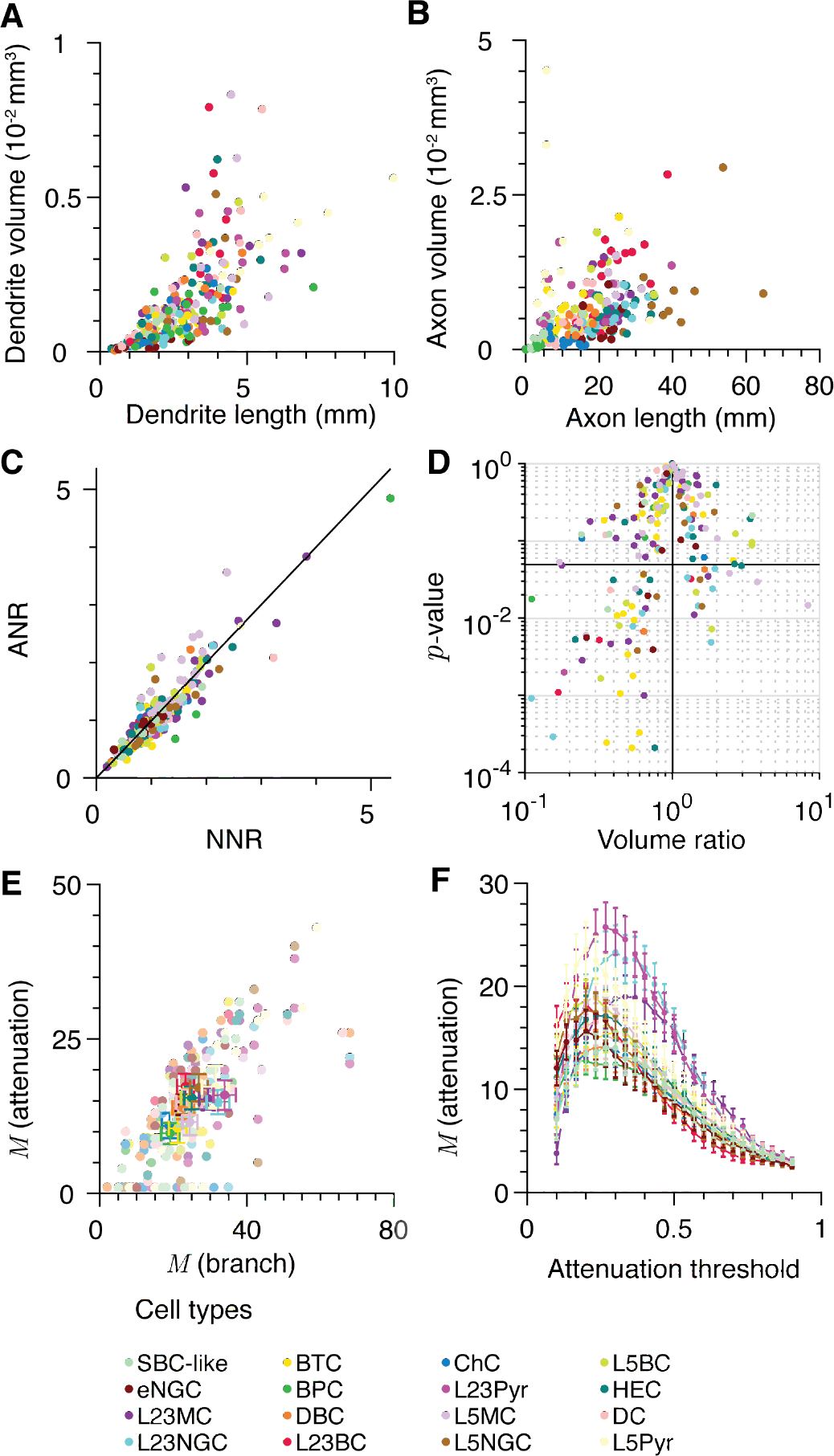
Supplement to Figure 4: Neurite size, neighbour ratios, and compartmentalisation for microcircuit data in Figure 4. **A** Dendrite spanning volume as a function of dendrite length for cells in the dataset. Colours indicate cell type (see legend). **B** As in **A**, for axons. **C** Relationship between nearest-neighbour ratio (NNR) and all-neighbour ratio (ANR) for each set of putative contacts. Colours indicate presynaptic cell type and the linear correlation coefficient is 0.91. **D** Volume ratio and p-value for sets of putative contacts. **E** Number of compartments estimated using the branch definition against the number using the attenuation definition. Individual cells are shown with transparency and the mean for each cell class by a solid colour. Horizontal and vertical error bars show the standard error in both definitions. **F** Number of compartments estimated using the attenuation definition for different cell classes as a function of the proportional attenuation threshold. Error bars show standard error.

**Fig S5.**
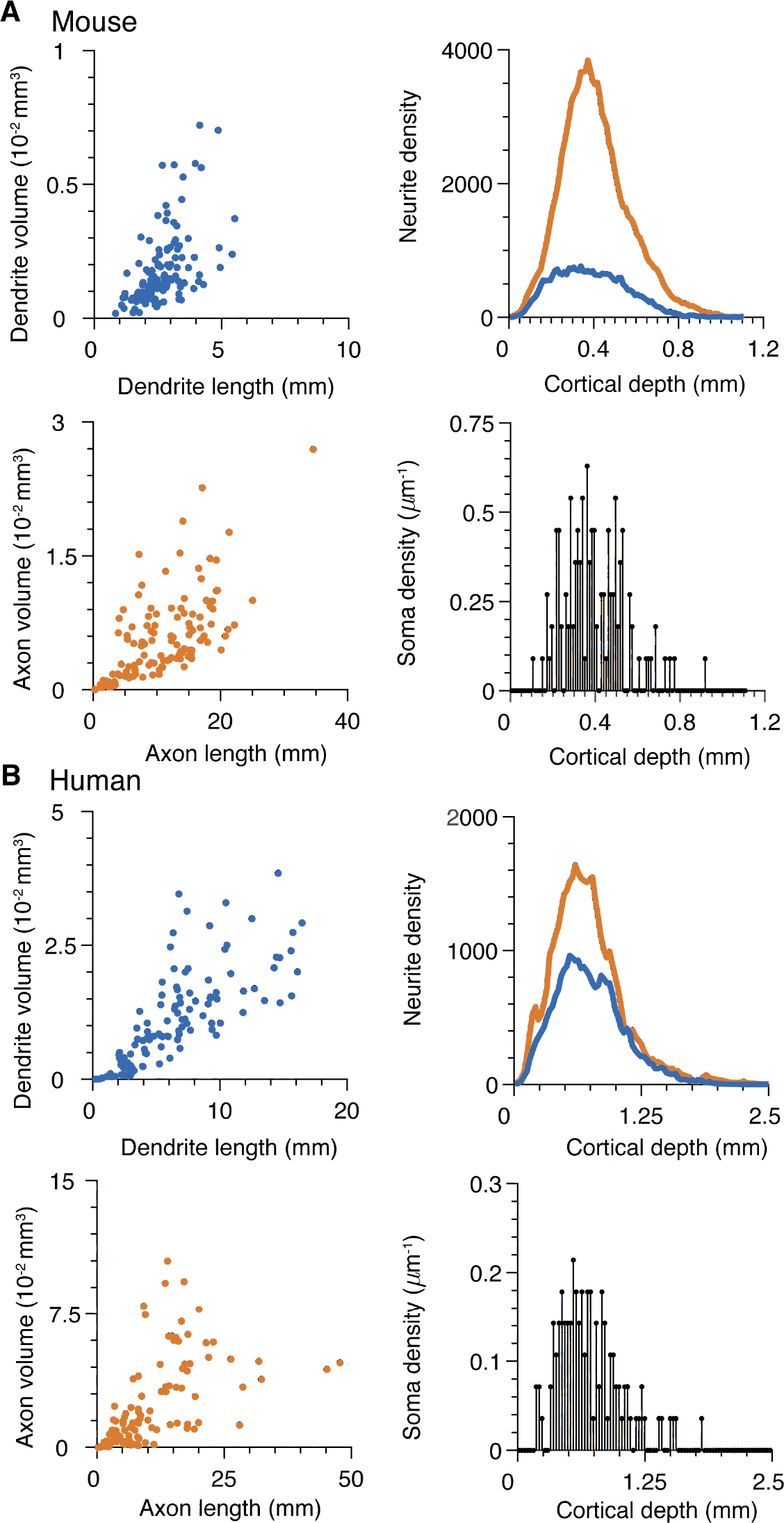
Supplement to Figure 5: Neurite size and cortical depth for the reconstruction data in Figure 5. **A** Mouse data. Top left: Dendrite spanning volume as a function of dendrite length for cells in the dataset. Bottom left: Axon spanning volume as a function of axon length. Top right: Neurite density (microns of neurite per *μ*m of cortical depth) for all axons (orange) and dendrites (blue) in the dataset. Bottom right: Soma density (soma per *μ*m of cortical depth) for all neurons in the dataset. **B** Human data. As in **A**, but for cortical human cells.

**Table S1.** Somatic input resistances and estimated membrane conductivities for the cell classes in Figure 4. Input resistance data taken from Jiang et al (2015), except for values labelled with * where Guan et al (2015) is the source. In all cases the axial resistance *ra* = 100 Ωcm.

| Cell type | Electrotonic properties |  |
| --- | --- | --- |
|  | Input resistance (soma) | Membrane conductivity |
| | (MΩ) | $g_m$ ($10^{-4}\text{Scm}^{-2}$) |
| SBC-like | $136.8 \pm 5.4$ | 1.941 |
| eNGC | $132.8 \pm 6.0$ | 2.155 |
| L23MC | $126.1 \pm 3.6$ | 1.216 |
| L23NGC | $120.6 \pm 7.1$ | 1.495 |
| BTC | $131.6 \pm 5.6$ | 1.389 |
| BPC | $131.2 \pm 5.4$ | 1.941 |
| DBC | $76.7 \pm 2.7$ | 2.765 |
| L23BC | $84.5 \pm 3.9$ | 1.941 |
| ChC | $111.7 \pm 5.8$ | 2.155 |
| L23Pyr | $120 \pm 10^*$ | 1.145 |
| L5MC | $141.8 \pm 7.1$ | 1.025 |
| L5NGC | $95.4 \pm 6.4$ | 2.155 |
| L5BC | $95.8 \pm 4.0$ | 1.766 |
| HEC | $111.5 \pm 4.9$ | 1.941 |
| DC | $141.5 \pm 15.6$ | 1.766 |
| L5Pyr | $130 \pm 7^*$ | 0.927 |

